# PNPLA3-I148M is a Neomorph that Interferes with Two Primary Hepatic Triglyceride Clearance Pathways

**DOI:** 10.1101/2024.10.29.620978

**Authors:** David J. Sherman, Lei Liu, Edward L. LaGory, Preston Fordstrom, Danielle Pruitt, Jose Barreda, Jiansong Xie, John Ferbas, Ingrid C. Rulifson, Raymond J. Deshaies

**Author notes:** Co-senior authors (Ingrid C. Rulifson, Raymond J. Deshaies). Correspondence: David J. Sherman, Raymond J. Deshaies.

## Abstract

A common variant of *PNPLA3*, encoding PNPLA3-I148M, is the most significant genetic determinant of fatty liver disease worldwide. However, it is unclear precisely how PNPLA3-I148M drives disease risk. Here, we show that endogenous human PNPLA3-I148M impairs secretion of Apolipoprotein B (ApoB), the scaffolding protein of very low-density lipoproteins (VLDLs), from hepatocytes. This is not due to a generalized secretory pathway defect, nor is it equivalent to loss of function of PNPLA3. The VLDL secretory defect is conserved in mice expressing human I148M. Untargeted lipidomics reveal that I148M human cells are enriched in polyunsaturated fatty acid (PUFA)-containing triglycerides at the expense of PUFA-containing phosphatidylcholine, causing reduced membrane dynamics, concomitantly hindering biogenesis of secreted VLDLs. ApoB secretion is substantially rescued in I148M cells that overexpress ABHD5/CGI-58, an I148M binding partner that activates lipolysis by ATGL/PNPLA2 when not bound to I148M. Conversely, knocking down CGI-58 or PNPLA2 mimics I148M. We propose that neomorphic PNPLA3-I148M exacerbates fatty liver risk by simultaneously impeding two major CGI-58-dependent pathways for liver triglyceride clearance: lipolysis and secretion.

## Main Text

A nonsynonymous variant in the gene *PNPLA3*, encoding an isoleucine-to-methionine change at position 148, is strongly associated with the full spectrum of metabolic dysfunction-associated steatotic liver disease (MASLD) worldwide^1–3^. Starting with hepatic steatosis, or excess triglyceride (TG) buildup in the liver, MASLD encompasses histological changes that include metabolic dysfunction-associated steatohepatitis (MASH), cirrhosis, and hepatocellular carcinoma^4^. As of 2023, MASLD was estimated to afflict ∼ 32% of the world adult population and is growing in prevalence in children and adults similarly to obesity and type 2 diabetes^5–7^. Steatosis results from an imbalance of TG uptake/synthesis and removal, but the complex web of lipid trafficking and metabolic pathways that contribute to disease development and progression remain poorly understood. Therefore, despite significant research efforts, there is a dearth of therapies to treat MASLD. Unlike many genetic variants that substantially increase disease risk, PNPLA3-I148M is remarkably common. For example, ∼ 50% of the Hispanic population are estimated to carry the disease-causing variant, which parallels the disproportionately high incidence of MASLD in this group^1,8,9^. Accordingly, a deeper understanding of the mechanism(s) by which PNPLA3-I148M enhances disease risk can greatly enhance efforts to find treatments for this silent epidemic.

Patatin-like phospholipase domain-containing protein 3 (PNPLA3) is primarily a lipid droplet (LD)- and membrane-associated protein that belongs to a conserved family of lipases found in organisms ranging from bacteria to humans^10–12^. Since the identification of its disease-causing variant in 2008, efforts have been made to uncover the function of wild-type PNPLA3 and to understand how a single isoleucine-to-methionine change increases risk of the full MASLD spectrum^12–19^. PNPLA3-I148M has been described both as a loss- and a gain-of-function variant, even though there is not a clear molecular understanding of the function of the wild-type protein in the broader context of human physiology. The traditional loss- or gain-of-function attributions fall short when considering that neither *Pnpla3* knock-out nor wild-type PNPLA3 overexpression in mice causes steatosis or phenotypes associated with MASLD^12,20–22^. However, it should be noted that murine Pnpla3 and human PNPLA3 differ in primary sequence and tissue distribution^23^.

There are two reported observations that provide insights into how PNPLA3-I148M might promote disease. One is the observation that PNPLA3-I148M binds to the protein α/β-hydrolase domain-containing protein 5 (ABHD5), also known as comparative gene identification-58 (CGI-58) ^15,16^. CGI-58 interacts with and activates the lipolytic activity of adipose triglyceride lipase (ATGL), also known as PNPLA2, the closest relative of PNPLA3^24,25^. PNPLA2 catalyzes the rate-limiting step of triglyceride hydrolysis, releasing diacylglycerol and a non-esterified fatty acid. Its enzymatic activity is increased up to ∼ 20-fold by CGI-58, although a molecular explanation of this activation remains elusive^24,26^. The interaction between PNPLA3-I148M and CGI-58 is thought to sequester CGI-58 from PNPLA2, effectively inhibiting its ability to catalyze the first step of triglyceride hydrolysis and leading to hepatic LD accumulation^27^. Naturally occurring loss-of-function *PNPLA2* and *CGI-58* variants have been identified in humans with neutral lipid storage diseases characterized by aberrant LD accumulation in different organs and tissues^2,28,29^. However, characteristics of the diseases caused by these variants differ, and mouse models with *Pnpla2* or *Cgi-58* deletions demonstrate distinct phenotypes that suggest less well-understood PNPLA2-independent roles for CGI-58 in regulating triglyceride metabolism^30^.

The second notable observation that provides clues as to how PNPLA3-I148M might cause MASLD is that individuals carrying this variant produce triglyceride-poor very low-density lipoprotein (VLDL) particles^17–19,31^. VLDLs are 35- to 100-nm diameter triglyceride- and phospholipid-rich particles coated with apolipoproteins, namely apolipoprotein B-100 (ApoB), a ∼ 550 kilodalton scaffolding protein found on every VLDL particle with a stoichiometry of one ApoB molecule per particle^32^. VLDL secretion is a major pathway for exporting hepatic triglycerides for energy use or storage elsewhere in the body^33,34^. VLDL particles are biosynthesized along the traditional secretory pathway in biochemically distinct vesicles due to their lipid-rich cargo^32^. Biosynthesis starts with the addition of core lipids, mainly triglycerides, to ApoB by microsomal triglyceride transfer protein (MTP) as ApoB is co-translationally inserted into the endoplasmic reticulum (ER) lumen^32,35^. ApoB levels, and therefore VLDL biosynthesis, are tightly regulated by lipid availability and hormones, and improperly lipidated ApoB is targeted for degradation^32,35,36^. Nascent VLDL particles are packaged into specialized secretory vesicles at the ER and transported to the Golgi apparatus, where they are further elaborated with lipids, mainly phospholipids, before packaging and trafficking to the plasma membrane for secretion^32,37,38^. Several biochemical and biophysical factors, including phosphatidylcholine biosynthesis and membrane fluidity, are required for proper biosynthesis of these lipid-rich secretory cargoes^39,40^. Notably, naturally occurring loss-of-function variants in *MTP* and *APOB* have been reported to cause hepatic steatosis^4,41,42^. Additionally, the second most significant genetic variant associated with MASLD, encoding a glutamate-to-lysine amino acid change in the protein TM6SF2, has been shown to reduce hepatic VLDL secretion^43,44^.

Given these disparate observations, we sought to uncover the mechanistic underpinnings of PNPLA3-I148M function using an endogenous human hepatoma system that we developed to investigate PNPLA3 and PNPLA3-I148M biology^11^. Coupled with studies using primary human hepatocytes and mouse models expressing human PNPLA3-I148M, we provide evidence that the PNPLA3-I148M•CGI-58 axis acts at the confluence of lipolysis and VLDL biogenesis to cause hepatic fat accumulation. These data support a model in which PNPLA3-I148M acts as a neomorph, gaining a function distinct from the wild-type protein.

## Results

### Endogenous human PNPLA3-I148M impairs ApoB secretion in hepatocytes

We previously showed that endogenous PNPLA3-I148M expression in Hep3B cells causes LD accumulation relative to wild-type (WT) cells and alters the cellular proteomic landscape in ways that resemble liver disease^11^. When characterizing the paired WT and I148M Hep3B cells, we found that oleic acid-treated I148M cells were deficient in secreting ApoB into the medium (Fig. 1a). Additionally, I148M cells lacked detectable intracellular ApoB by immunoblotting (Fig. 1a). To determine if this phenotype resembles loss-of-function of WT PNPLA3, we generated *PNPLA3* knockout cells and measured ApoB secretion into the media (Fig. S1). The *PNPLA3* knockout clones resembled WT Hep3B cells, without a significant difference in ApoB levels in the media, as measured by an ApoB Enzyme Linked Immunosorbent Assay (ELISA; Fig. 1b).

**Figure 1:**
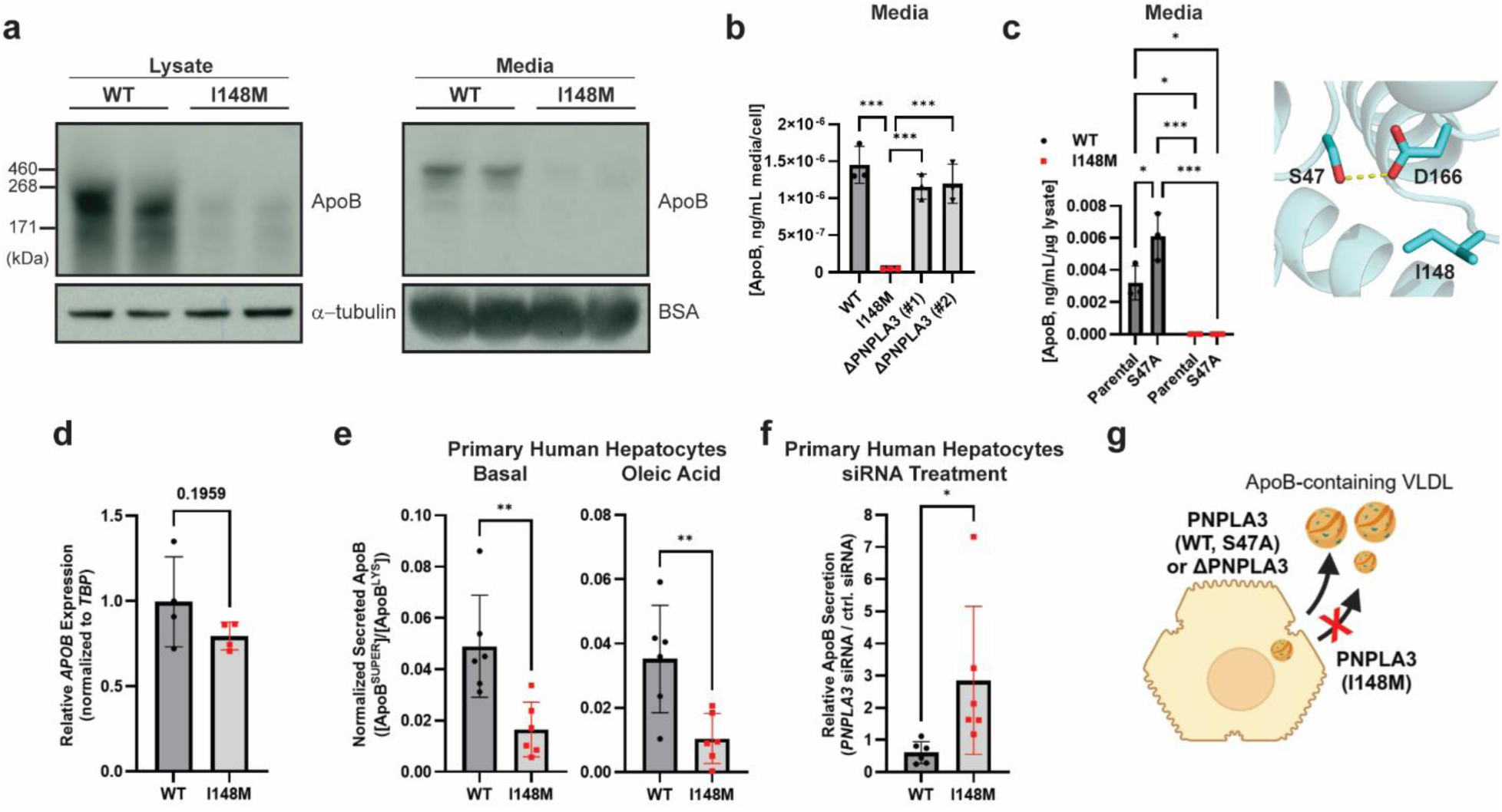
PNPLA3-I148M interferes with ApoB secretion in human hepatocytes. **a,** Representative ApoB immunoblots of lysate and media from Hep3B cells. **b,** Quantification of ApoB in the media of oleic acid (100 µM)-treated Hep3B cells (WT, I148M, and two PNPLA3 knockout clones), measured by ELISA. Analyzed by ordinary one-way Anova with Tukey’s multiple comparisons test. WT-I148M, adj. *p* = 0.0001; I148M-ΔPNPLA3 #1, adj. *p* = 0.0006; I148M-ΔPNPLA3 #2, adj. *p* = 0.0005. *n* = 3 biological replicates. **c,** Quantification of ApoB in the media of oleic acid (100 µM)-treated Hep3B cells (WT, WT_S47A, I148M, I148M_S47A), measured by ELISA. Analyzed by two-way Anova with Tukey’s multiple comparisons test. WT-WT_S47A, adj. *p* = 0.0186; WT-I148M, adj. *p =* 0.0107; WT-I148M_S47A, adj. *p* = 0.0107; I148M-WT_S47A, adj. *p* = 0.0002; WT_S47A-I148M_S47A, adj. *p* = 0.0002. *n* = 3 biological replicates. Zoomed-in active site of predicted PNPLA3 structure depicted on the right with catalytic dyad and I148 side chains illustrated as sticks (Alphafold, Uniprot Q9NST1). Carbon and oxygen are depicted in cyan and red, respectively. **d,** qPCR of RNA isolated from WT and I148M Hep3B cells. *APOB* levels were normalized to the those of the housekeeping gene *TPB*. Analyzed by unpaired *t-*test. WT-I148M, adj. *p* = 0.1959. *n* = 3 biological replicates. **e,** Lots of primary human hepatocytes (*n =* 3 and *n =* 4 lots for WT and I148M, respectively) were incubated without (left) or with (right) 200 µM oleic acid. ApoB in the lysate and the media were quantified by ApoB ELISA and the ratio of ApoB in the media supernatant (“SUPER”) to lysate (“LYSATE”) is shown. Analyzed by unpaired *t-* test, adj. *p* = 0.0055 (basal) and *p =* 0.0081 (oleic acid treatment). *n* = 6 biological replicates. **f,** Two lots of WT and two lots of I148M primary human hepatocytes were treated with 1 nM control siRNA or PNPLA3-targeting siRNA, followed by addition of 200 µM oleic acid prior to analyzing ApoB content in the lysate and media by ELISA. Analyzed by unpaired *t-*test, adj. *p* = 0.0055 (basal) and *p =* 0.0398. *n* = 6 biological replicates. **g,** Schematic summarizing the consequences on ApoB secretion of various forms of PNPLA3.^†^ * *p* ≤ 0.05; ** *p* ≤ 0.01; *** *p* ≤ 0.001.

Because the *PNPLA3* knockout clones phenocopied WT Hep3B cells, we next asked whether introducing a point mutation in the putative PNPLA3 active site affects ApoB secretion. Serine 47 is a part of the conserved catalytic dyad found in patatin-like phospholipase domain-containing proteins^10^. Therefore, we knocked-in the serine-47-alanine (S47A) variant at the endogenous locus in WT and I148M cells (Fig. S2). PNPLA3-S47A has previously been shown to inhibit PNPLA3-mediated triglyceride lipase activity in a purified enzyme assay^13^, and unlike WT PNPLA3 is unable to sustain LD hydrolysis when expressed in *PNPLA2* knockout HeLa cells^19^. The S47A-I148M double mutant was indistinguishable from I148M in terms of ApoB secretion, suggesting that I148M function is not due to a gain of catalytic activity at the conserved active site (Fig. 1c). The S47A change in the WT Hep3B background did not hinder ApoB secretion, and in fact increased secretion, suggesting that the secretory defect is not due to a loss of enzymatic activity of the WT protein (Fig. 1c).

We confirmed that the loss of intracellular and secreted ApoB in I148M cells was not due to reduced *APOB* gene expression, as *APOB* mRNA levels were similar in WT and I148M cells (Fig. 1d). This result suggests that a co- or post-translational degradation mechanism, or possibly a change in the rate of protein synthesis, reduces ApoB protein levels. ApoB can be targeted for degradation by both the ubiquitin-proteasome system and autophagic pathways^36^. To test whether the low level of ApoB in I148M cells is due to its degradation, we treated Hep3B cells with CB-5083, a potent inhibitor of the valosin-containing protein (VCP)/p97, which is involved in both ER-associated degradation by the proteasome and autophagy^45,46^. As expected, CB-5083 treatment caused accumulation of polyubiquitin conjugates and the ER-targeted transcription factor NFE2L1/Nrf1, a protein that is constitutively degraded in a p97-dependent pathway unless p97 and/or the ubiquitin-proteasome system are inhibited (Fig. S3a) ^47^. Notably, p97 inhibition caused significant intracellular accumulation of ApoB in both WT and I148M cells (Fig. S3b), consistent with the observation that a significant pool of ApoB that is synthesized is destined for degradation^36^. This result suggests that low homeostatic levels of ApoB in I148M cells are maintained by protein quality control pathways.

We next tested whether endogenous PNPLA3-I148M inhibits ApoB secretion in primary human hepatocytes. We identified primary hepatocytes from donors expressing WT *PNPLA3* and donors homozygous for *PNPLA3-*I148M (Table S1). Unlike in Hep3B cells, the primary hepatocytes had comparable levels of intracellular ApoB across primary cell lots (i.e., from different donors), but the levels of secreted ApoB differed by *PNPLA3* status. Therefore, as a measure of secreted ApoB, levels of ApoB in the supernatant/medium (“SUPER”) were normalized to levels in the lysate (“LYSATE”). Using the ratio instead of absolute ApoB in the medium also provided a measure to control for differences in housekeeping gene expression across donors. Both with and without oleic acid stimulation, the WT lots secreted more ApoB than the I148M lots (Fig. 1e). Given that the lots came from different individuals, we wanted to confirm that the observed effect was due to expression of PNPLA3-I148M and not due to other genetic differences. Therefore, we treated WT and I148M primary hepatocytes (two lots per group) with small interfering RNA (siRNA) targeting PNPLA3. Knockdown was highly efficient in all primary cells tested (> 80% knockdown, as measured by digital PCR; data not shown). ApoB secretion was increased ∼ 3-fold upon knockdown of PNPLA3-I148M in primary hepatocytes, which was not observed for WT PNPLA3 knockdown (Fig. 1f). Cumulatively, these data suggest that the defect is specific to I148M and is not caused by loss of PNPLA3 enzymatic activity or loss of PNPLA3 protein (Fig. 1g).

### Human PNPLA3-I148M reduces VLDL triglyceride levels in mice without a dietary challenge

ApoB is the main scaffolding protein for VLDL particles, which are rich in TG content. Therefore, we sought to determine if the ApoB secretory defect caused by PNPLA3-I148M in cells translates to reduced VLDL-TG secretion *in vivo*. We generated mice with the full human PNPLA3-I148M coding sequence knocked-in at the endogenous mouse locus, replacing the mouse *Pnpla3* gene. The cohorts of mice were phenotypically indistinguishable despite the differences at the *Pnpla3* locus (Fig. S4a). Following four weeks on a CHOW diet, which does not induce MASLD phenotypes on its own, WT and huPNPLA3-I148M knock-in (KI; both heterozygous, HET, and homozygous, HOM) mice were fasted overnight. Plasma from each group was pooled and fractionated to separate lipoproteins based on size (Fig. 2a). huPNPLA3-I148M expression in mice on a CHOW diet for the duration of the study did not cause an increase in liver size relative to body size (in fact, the liver weight to body weight ratio of HOM mice was reduced relative to WT and HET mice; Fig. 2b). huPNPLA3-I148M expression did not cause liver TG accumulation, but the HOM mice had slight accumulation of hepatic cholesterol (Chol; Fig. 2c). However, the VLDL fractions of huPNPLA3-I148M KI mice contained less TG than their WT counterparts, following a huPNPLA3-I148M dose response (HET and HOM VLDL-TG values were ∼ 29% and ∼ 50%, respectively, less than WT, as judged by the area under the curve; Fig. 2d). HOM mice also had less cholesterol in their high-density lipoprotein (HDL) fractions (Fig. 2d). Although humans secrete full-length ApoB100 from the liver, the majority of ApoB secreted from the mouse liver is an mRNA-edited version called ApoB48, which represents the first 48% of the ApoB100 protein^48,49^. VLDL ApoB levels, as measured by an ELISA that recognizes both ApoB48 and ApoB100 forms, followed a similar trend as the TG levels (Fig. 2e).

**Figure 2:**
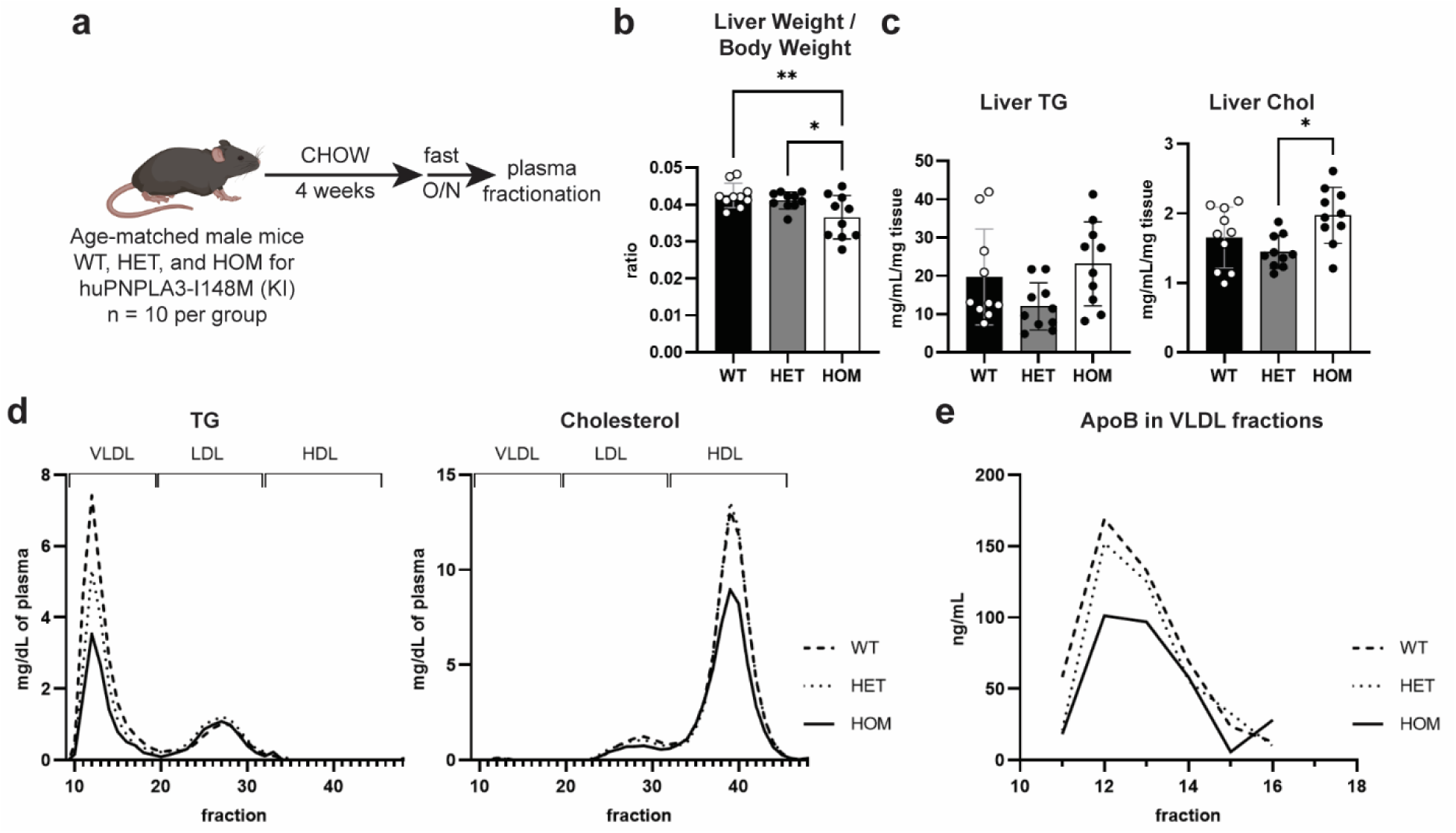
Human PNPLA3-I148M expression reduces VLDL-TG secretion in mice. **a,** Schematic diagram of huPNPLA3-I148M knock-in mouse study on the CHOW diet.^†^ **b,** Liver weight-to-body weight ratio of mice. Analyzed by ordinary one-way Anova with Tukey’s multiple comparisons test. WT-HOM, adj. *p* = 0.0086; HET-HOM, adj. *p* = 0.0486. **c,** Quantification of liver triglyceride and liver cholesterol content. Analyzed by ordinary one-way Anova with Tukey’s multiple comparisons test. Liver Chol: HET-HOM, adj. *p* = 0.0103. **d,** Pooled mouse plasma following an overnight fast was subject to size-exclusion chromatography on a Superose 6 column and TG and Chol levels were quantified in the fractions. **e,** VLDL fractions were analyzed by ApoB ELISA. * *p* ≤ 0.05; ** *p* ≤ 0.01.

To obtain improved resolution of the sizes of the VLDL particles secreted by the mice, pooled plasma from CHOW-fed WT and HOM mice was subjected to size-exclusion chromatography on a Sephacryl S-500 column, which is better able to resolve larger-sized particles. In the resulting chromatograms, the HOM VLDL-TG curve was right-shifted relative to the WT curve (Fig. S4b). Because larger VLDL particles elute in the earlier (left-most) fractions, this result suggests that the VLDL particles secreted by HOM mice are smaller in size than those of WT mice. The three groups (WT, HET, and HOM) had similar liver expression profiles of various lipid metabolic and VLDL biogenesis genes (Fig. S4c). Hence, we concluded that endogenous expression of huPNPLA3-I148M in CHOW-fed mice impairs the biogenesis of plasma VLDL.

Two groups of mice (WT and HOM KI) were also fed the American lifestyle-induced obesity syndrome (ALIOS) diet, which has been reported to induce the full MASLD spectrum over extended periods of time^50^. Following 3 weeks on the ALIOS diet, which is too short to cause MASLD, mice were fasted overnight, plasma was pooled by group and fractionated to separate lipoprotein particles (Fig. S5a). A parallel group of CHOW-fed HOM KI mice was included in this study for comparison. There were insignificant changes in liver weight to body weight ratios and hepatic TG levels on the ALIOS diet, although the HOM ALIOS group had some Chol accumulation relative to their WT counterparts (Fig. S5b,c). The groups followed the same VLDL secretion trend as on the CHOW diet, with HOM huPNPLA3-I148M KI mice exhibiting reduced VLDL secretion relative to WT mice (Fig. S5d,e). Hepatic expression changes of various lipid metabolic and VLDL biosynthetic genes could not explain the genotype-specific differences on the ALIOS diet (Fig. S5f). These results provide further support for a model in which huPNPLA3-I148M impairs hepatic VLDL-TG biogenesis and secretion in mice.

### The effect of PNPLA3-I148M on ApoB secretion is neither due to a generalized secretory defect nor LD accumulation

With evidence that the cellular ApoB secretory defect translates to altered VLDL biogenesis and secretion *in vivo*, we next considered the possibility that I148M cells have a general secretory defect. Like traditional secretory cargoes, nascent VLDL particles are trafficked from the ER to the Golgi, although the VLDL trafficking process involves additional protein machinery and proteins that add lipids to the growing particles^32^. To probe whether PNPLA3-I148M impairs the general secretory pathway, cells were transfected with a vector to express *Gaussia* luciferase, a small (20 kDa), soluble secretory protein that generates luminescence in the presence of the substrate coelenterazine and oxygen^51^. Following a media exchange, cells were treated with or without Brefeldin A (BFA), an inhibitor of ER-to-Golgi secretory protein trafficking^52^, and luminescence was measured in the conditioned media to determine *Gaussia* luciferase secretion after 24 h (Fig. 3a). Both WT and I148M cells secreted *Gaussia* luciferase, with I148M cells secreting more than WT cells (Fig. 3a). Treatment of either cell type with BFA, or lack of *Gaussia* luciferase expression, reduced or eliminated luminescence in the media. Therefore, I148M cells are effectively able to secrete a small, soluble protein substrate. In addition to the *Gaussia* luciferase functional assay, immunoblotting for various components of the traditional secretory pathway indicated similar protein expression levels in WT and I148M cells (Fig. 3b). Therefore, I148M cells are not deficient in secretory pathway components.

**Figure 3:**
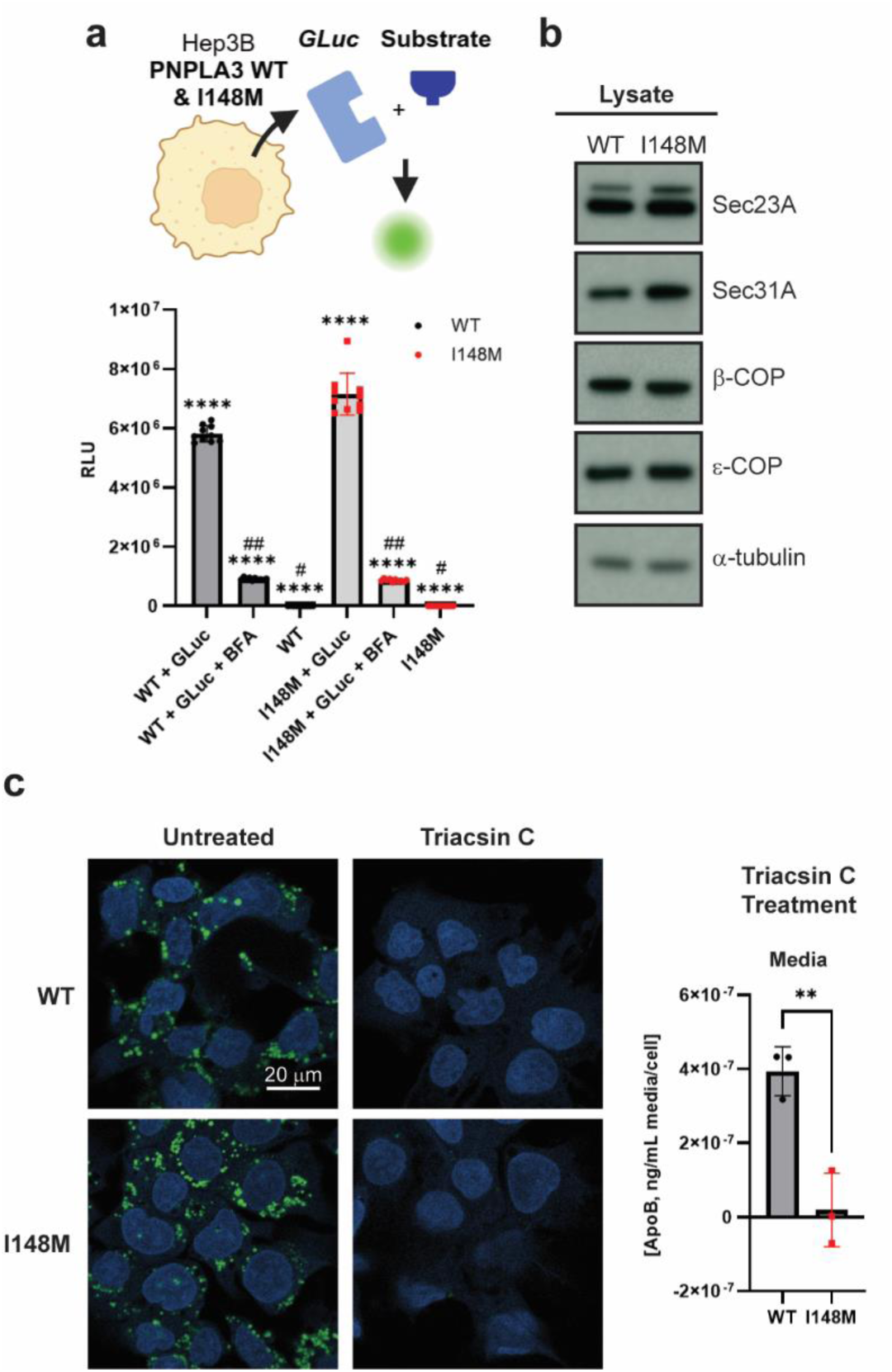
The VLDL secretory defect caused by PNPLA3-I148M is not a generalized secretory defect and is independent of cytosolic lipid droplet content. **a,** Hep3B cells were transfected with a vector to express the secreted soluble protein *Gaussia* luciferase. Some samples were additionally treated with Brefeldin A (5 µg/mL) for 24 h to inhibit the secretory pathway. Media was removed and incubated with coelenterazine to measure the luminescence generated by secreted luciferase. Analyzed by ordinary one-way Anova with Tukey’s multiple comparisons test. **** adj. *p* < 0.0001 compared to every other sample unless otherwise noted; # adj. *p* > 0.05 compared to each other; ## adj. *p* > 0.05 compared to each other. *n =* 10 replicates, repeated twice.^†^ **b,** Representative immunoblots of various secretory pathway components in the lysates of WT and I148M Hep3B cells. **c,** Hep3B cells were treated with Triacsin C (5 µM) for 24 h to clear lipid droplets from cells. Media was then exchanged for fresh media containing Triacsin C (5 µM) and oleic acid (100 µM), and cells were incubated for an additional 24 h prior to analyzing ApoB content in the media by ELISA. Some cells were fixed after 24 h with Triacsin C, and lipid droplets and nuclei were stained with LipidTOX Green and Hoechst 33342 (blue), respectively (left). ApoB in the media from cells treated with Triacsin C (48 h total) and oleic acid (24 h) was quantified by ELISA (right). Analyzed by unpaired *t-*test, adj. *p* = 0.0056. *n* = 3 biological replicates. ** *p* ≤ 0.01.

Although we reasoned that failure to secrete ApoB-containing lipoproteins could cause intracellular TG accumulation, we also considered the possibility that higher steady-state levels of LDs in I148M cells could somehow impair VLDL biogenesis. To probe this “chicken and egg” question, WT and I148M cells were treated with Triacsin C, an inhibitor of long-chain acyl-coenzyme A (CoA) synthetase that triggers LD breakdown. After effectively clearing the WT and I148M cells of LDs (Fig. 3c), cells were stimulated with oleic acid to promote VLDL biosynthesis. Clearing the cells of LDs did not rescue the ApoB secretory defect in I148M cells (Fig. 3c). Therefore, LD accumulation is likely a consequence, not a cause, of the ApoB secretory defect.

### I148M cells have altered phosphatidylcholine composition and reduced membrane dynamics

Membrane phospholipid composition can affect VLDL biogenesis^39,40,53^. Therefore, we next asked whether differences in lipid composition in WT and I148M cells could explain the ApoB secretory defect. WT and I148M cells were treated with or without oleic acid for 24 h prior to harvesting the cells and extracting lipids for characterization by liquid chromatography-mass spectrometry (LC-MS). Untargeted lipidomics analysis revealed significant differences in lipid composition between the WT and I148M cells: 230 and 402 lipid species differed between groups in basal and oleic acid-treated conditions, respectively (Fig. 4a). TG and cholesterol esters were especially elevated in I148M cells. Notably, there were deficiencies in various PC species in I148M cells, especially with oleic acid treatment. However, total PC content did not differ between groups (Fig. 4b).

**Figure 4:**
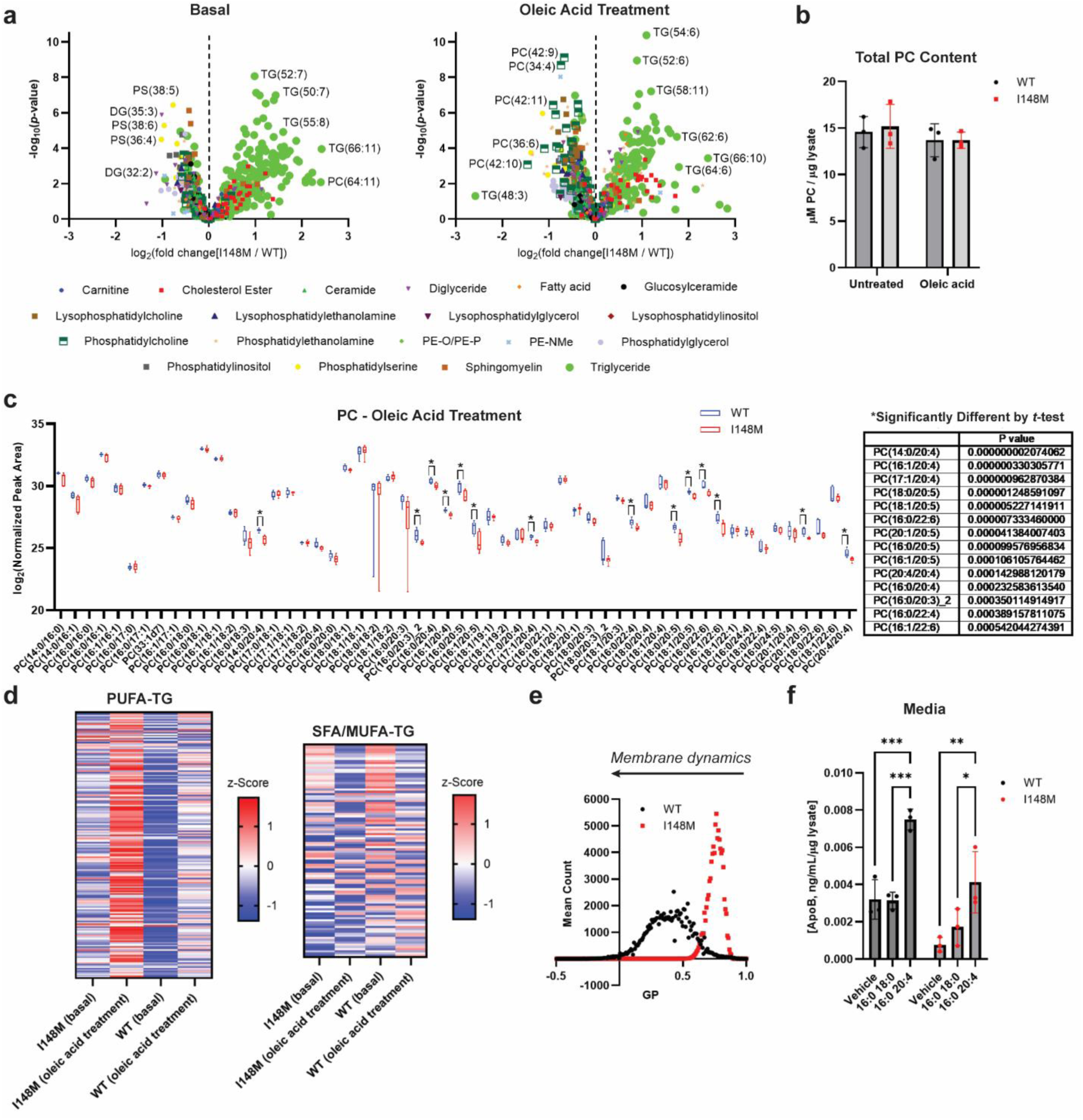
PNPLA3-I148M cells have reduced polyunsaturated fatty acid (PUFA)-phosphatidylcholine (PC) content and impaired membrane dynamics. **a,** Volcano plots of lipid species from basal (untreated; left) or oleic acid-treated (100 µM for 24 h; right) Hep3B cells (WT and I148M). Certain significantly changing lipids are annotated. *n* = 3 biological replicates. Parenthetical lipid designations refer to (Total # of carbons across acyl chains : Total # of double bonds across acyl chains). **b,** Total cellular phosphatidylcholine content was measured in untreated or oleic acid (100 µM)-treated Hep3B WT and I148M cells. Analyzed by two-way Anova with Tukey’s multiple comparisons test. All adj. *p* > 0.7. *n* = 3 biological replicates. **c,** Plot of normalized peak area for detected PC species in oleic acid-treated cells (left). Analyzed by *t-*tests with the Holm-Sidak method to correct for multiple comparisons. All significant adj. *p*-values (*p* < 0.05) given in the table (right). **d,** Heat maps of PUFA-TG and saturated fatty acid/monounsaturated fatty acid (SFA/MUFA)-TG species in untreated and treated Hep3B WT and I148M cells. **e,** Hep3B cells were incubated with the polarity-sensitive membrane dye di-4-ANEPPDHQ to measure membrane dynamics. Mean generalized polarization (GP) was calculated from live-cell confocal fluorescence microscope images. 80-90 cells were imaged for each sample (WT and I148M). **f,** Hep3B cells were treated for 24 h with oleic acid (100 µM) supplemented with vehicle (PBS) or liposomes (100 µM) of PC(16:0/18:0) or PC(16:0/20:4). ApoB content in the media was measured by ELISA. Analyzed by two-way Anova with Tukey’s multiple comparisons test. WT: vehicle-PC(16:0/20:4), adj. *p* = 0.0004; PC(16:0/18:0)-PC(16:0/20:4), adj. *p* = 0.0003. I148M: vehicle-PC(16:0/20:4), adj. *p* = 0.0028; PC(16:0/18:0)-PC(16:0/20:4), adj. *p* = 0.0238. *n =* 3 biological replicates. For **f:** * *p* ≤ 0.05; ** *p* ≤ 0.01; *** *p* ≤ 0.001.

Quantifying the normalized peak areas for PC species revealed that I148M cells were deficient in PCs containing polyunsaturated fatty acids (PUFAs; Fig. 4c). While one PC-PUFA species (PC 16:1 20:5) was significantly (*p*-value < 0.0005) decreased in I148M cells relative to WT cells under basal conditions, fourteen PC-PUFAs were reduced in I148M cells with oleic acid treatment using this same *p*-value threshold (Fig. 4c). Because absolute PC levels did not differ between the groups, we reasoned that either I148M cells had reduced PUFAs, or the PUFAs were sequestered in non-PC phospholipids or TG. In fact, TGs in I148M cells were enriched in PUFAs – especially with oleic acid treatment – whereas WT TGs were enriched in saturated fatty acids (SFAs) and monounsaturated fatty acids (MUFAs; Fig. 4d). This is a similar trend to what has been observed in the livers of humans homozygous for *PNPLA3*-I148M^18^.

PC is the major phospholipid found in secreted VLDL particles, as well as the predominant phospholipid component of cellular membranes, and impaired PC biosynthesis has been shown to hinder VLDL biosynthesis^39^. However, we did not observe global deficiencies in PC levels (Fig. 4b). Therefore, if PC composition in I148M causes the ApoB secretory defect, we reasoned that it might be due to the biophysical properties of the membranes involved in making VLDL particles. PUFA-containing phospholipids influence properties such as membrane fluidity, packing, thickness, and rigidity, which are important parameters to enable the large deformations and curvature required to make lipoproteins^40,54–56^. We interrogated whether a decrease in PC-PUFAs in I148M cells affected membrane dynamics by treating cells with the polarity-sensitive membrane dye di-4-ANEPPDHQ^57,58^. This permeable fluorophore partitions into cellular membranes and exhibits a 60-nm spectral shift in its emission between disordered and ordered membrane phases. Therefore, this experiment allowed us to quantify the degree of intracellular membrane order, a reflection of the dynamic nature of the membrane phospholipids, through a ratiometric fluorescence intensity measure known as generalized polarization (GP) ^59^. I148M cells exhibited a higher mean GP value for their intracellular membranes than WT cells (Fig. 4e), indicating that they have a greater degree of membrane order and reduced membrane dynamics. WT cells had a broader distribution of GP values, shown by the histogram in Fig. 4e.

To test whether the ApoB secretory defect in I148M cells is due to the impaired membrane dynamics observed with di-4-ANEPPDHQ, WT and I148M cells were treated with PC liposomes containing a saturated fatty acid (stearic acid) or a PUFA (arachidonic acid) at the *sn*-2 position. Supplementation with a PC-PUFA rescued ApoB secretion in I148M cells to basal WT levels (Fig. 4f). The fact that ApoB secretion also increases in WT cells supplemented with PC-PUFA suggests membrane fluidity can be further augmented to support ApoB secretion in the context of WT PNPLA3. Cumulatively, these results support a model in which I148M cells have impaired VLDL biogenesis because of their higher membrane order (higher GP), a result of lower PC-PUFA levels.

### The PNPLA3-I148M-induced ApoB secretory defect is mediated by CGI-58

We next sought to determine specifically how PNPLA3-I148M expression causes the lipidomic changes that impair VLDL biogenesis. Because the data do not support a model in which I148M possesses an enzymatic gain or loss of function (Fig. 1b,c), we asked whether the reported protein-protein interaction between PNPLA3-I148M and CGI-58 is responsible for this phenotype. First, we sought to confirm that we could observe decreased enzymatic lipolysis in I148M cells, as would be expected if PNPLA3-I148M sequesters CGI-58. Hep3B cells were starved for four hours to induce TG breakdown, and LDs were imaged and quantified during the time course. Some samples contained Bafilomycin A1, an inhibitor of autophagy, which would block lipophagic degradation of LDs, an alternative to enzymatic lipolysis^60^. WT cells followed a similar trend during the time course, with minimal difference between the samples treated with and without Bafilomycin A1 (Fig. S6). However, Bafilomycin A1 blocked LD breakdown in I148M cells (Fig. S6), consistent with reduced enzymatic lipolysis and thus a greater dependence on lipophagy. This response to autophagy inhibition supports a model in which PNPLA3-I148M sequesters CGI-58 in the Hep3B cells, inhibiting enzymatic lipolysis of TG.

To see whether the ApoB secretory defect is influenced by the interaction of PNPLA3-I148M with CGI-58, CGI-58 was knocked-down in WT and I148M cells by two independent siRNA molecules targeting different parts of the protein coding region, resulting in undetectable CGI-58 protein levels, as judged by immunoblotting (Fig. 5a). CGI-58 knockdown reduced ApoB secretion in WT cells by ∼ 75%, while ApoB secretion in I148M cells was similar to control siRNA levels (Fig. 5b). Next, we wondered how an excess of CGI-58, such that more CGI-58 is unbound by PNPLA3-I148M, would affect ApoB secretion in I148M cells. We generated WT and I148M cells stably overexpressing CGI-58 with a C-terminal FLAG tag (CGI-58-FLAG) and observed a marked reduction in intracellular LD levels in these cell lines, relative to parental cells (Fig. 5c). Oleic acid treatment to stimulate ApoB secretion caused some accumulation of LDs in these cells, but the total LD dry mass was reduced to ∼ 50% that of the parental cells (Fig. 5c). Under these conditions, CGI-58 overexpression rescued the ApoB secretory defect in I148M cells, boosting secretion to basal WT levels (Fig. 5d,e). Additionally, CGI-58 overexpression doubled ApoB secretion in WT cells, suggesting a general role for CGI-58 in VLDL biogenesis, as has been reported in cellular systems and mouse models^61–63^. This result suggests that the PNPLA3-I148M-induced ApoB secretory defect is mediated at least in part by its interaction with CGI-58 and can be substantially rescued by overexpressing CGI-58.

**Figure 5:**
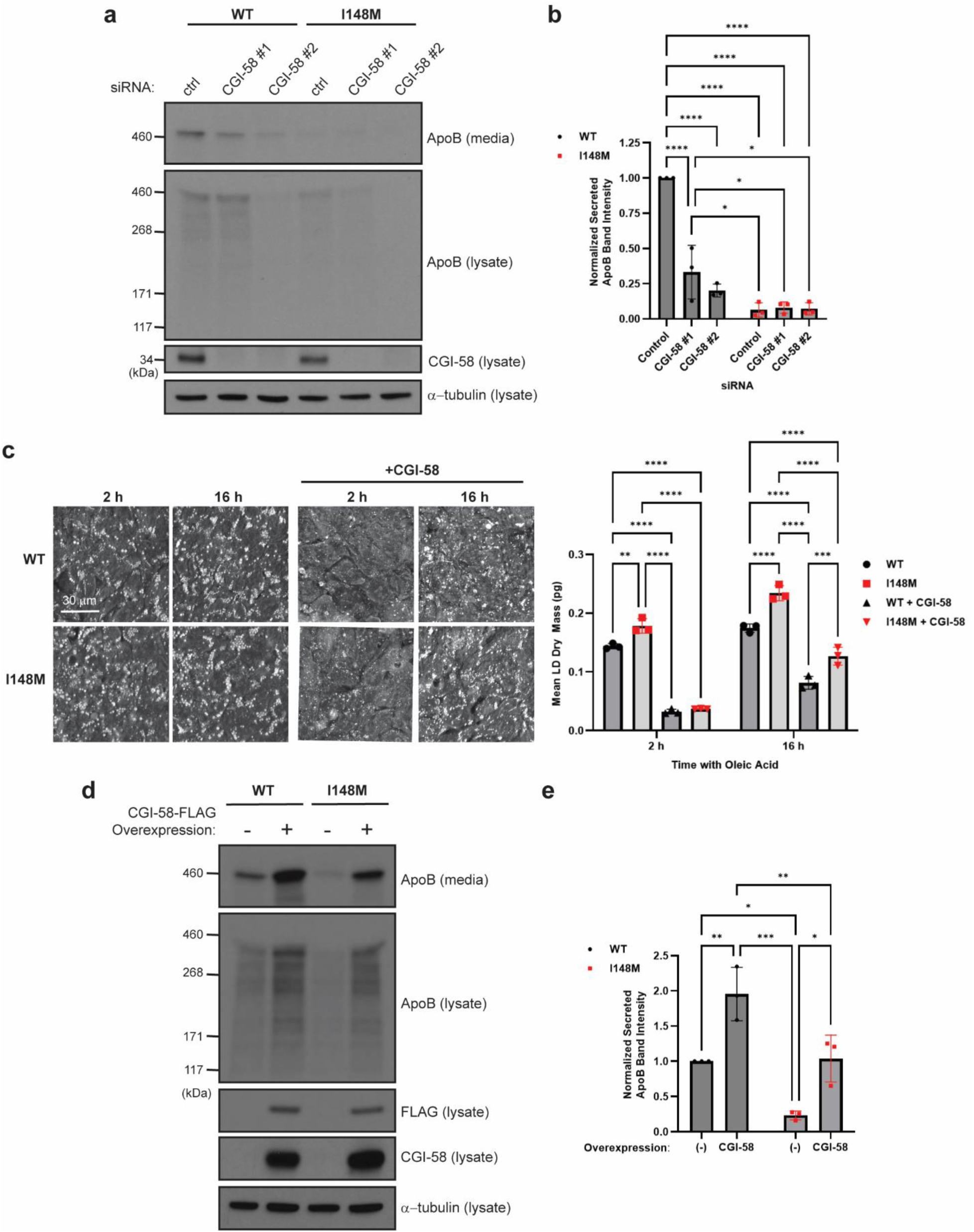
The ApoB secretory defect caused by PNPLA3-I148M is mediated by CGI-58. **a,**Hep3B cells were treated for 72 h with control (non-targeting) or CGI-58-targeting siRNA molecules. For the final 24 h, the media was exchanged for oleic acid (100 µM)-containing media. ApoB levels in the lysate and media were determined by immunoblots. **b,** ApoB immunoblots (**a**) were quantified by densitometry to determine relative ApoB levels in the media. Analyzed by two-way Anova with Tukey’s multiple comparisons test. WT control-I148M control, WT control-WT CGI-58 #1, WT control-WT CGI-58 #2, WT control-I148M CGI-58 #1, WT control-I148M CGI-58 #2 all have adj. *p* < 0.0001; I148M control-WT CGI-58 #1, adj. *p* = 0.0233; WT CGI-58 #1-I148M CGI-58 #1, adj. *p* = 0.0342; WT CGI-58 #1-I148M CGI-58 #2, adj. *p* = 0.0283. *n =* 3 biological replicates. **c,** Parental Hep3B cells and Hep3B cells stably overexpressing CGI-58-FLAG were treated with oleic acid (200 µM) for 16 h and imaged every ten minutes (starting 2 h after oleic acid addition) using Nanolive, a 3D holotomographic microscope that allows for lipid droplet identification by their refractive index (left). Mean lipid droplet dry mass (pg) in the time course was quantified (right). Analyzed by two-way Anova with Tukey’s multiple comparisons test. 2 h: WT-I148M, adj. *p* = 0.0026; WT-WT + CGI-58, WT-I148M + CGI-58, I148M-WT + CGI-58, I148M-I148M + CGI=58, adj. *p* < 0.0001. 16 h: all comparisons, except WT + CGI-58-I148M + CGI-58, adj. *p* < 0.0001. n *=* 3 biological replicates. **d,** Parental Hep3B cells and Hep3B cells stably overexpressing CGI-58-FLAG were treated as in **a**, and ApoB was detected in the lysate and media by immunoblotting. **e,** ApoB immunoblots (**d**) were quantified by densitometry to determine relative ApoB levels in the media. Analyzed by two-way Anova with Tukey’s multiple comparisons test. WT-I148M, adj. *p* = 0.0252; WT-WT + CGI-58, adj. *p* = 0.0078, I148M-WT + CGI-58, adj. *p* = 0.0002; I148M-I148M + CGI-58, adj. *p* = 0.0199; WT + CGI-58-I148M + CGI-58, adj. *p* = 0.0098. *n* = 3 biological replicates. * *p* ≤ 0.05; ** *p* ≤ 0.01; *** *p* ≤ 0.001; **** *p* ≤ 0.0001.

Next, we asked if the mechanism by which interaction of PNPLA3-I148M with CGI-58 inhibits ApoB secretion is related to inhibition of lipolysis by PNPLA2. We reasoned that lipolysis by PNPLA2 could release PUFAs from TG for incorporation into PC, facilitating membrane dynamics and VLDL biogenesis. PNPLA2 was knocked-down in WT and I148M cells by two independent siRNA molecules targeting different portions of the protein coding region. PNPLA2 protein levels were undetectable following knockdown, as judged by immunoblotting (Fig. 6a). PNPLA2 knockdown reduced ApoB secretion in WT cells by ∼ 60% (Fig. 6b). ApoB secretion in I148M cells was similar between the control and PNPLA2-targeting siRNA treatment groups. We next generated WT and I148M cells stably overexpressing PNPLA2 and found that overexpression boosted ApoB secretion in I148M cells to WT levels but did not significantly change WT ApoB secretion levels (Fig. 6c,d). These data support a model in which PNPLA3-I148M impairs ApoB secretion through its interaction with CGI-58, effectively mimicking loss of function of CGI-58. The PNPLA3-I148M•CGI-58 interaction inhibits lipolysis by PNPLA2 to free PUFAs from TG for incorporation into PC molecules, hindering membrane dynamics and impeding VLDL biogenesis (Fig. 7).

**Figure 6:**
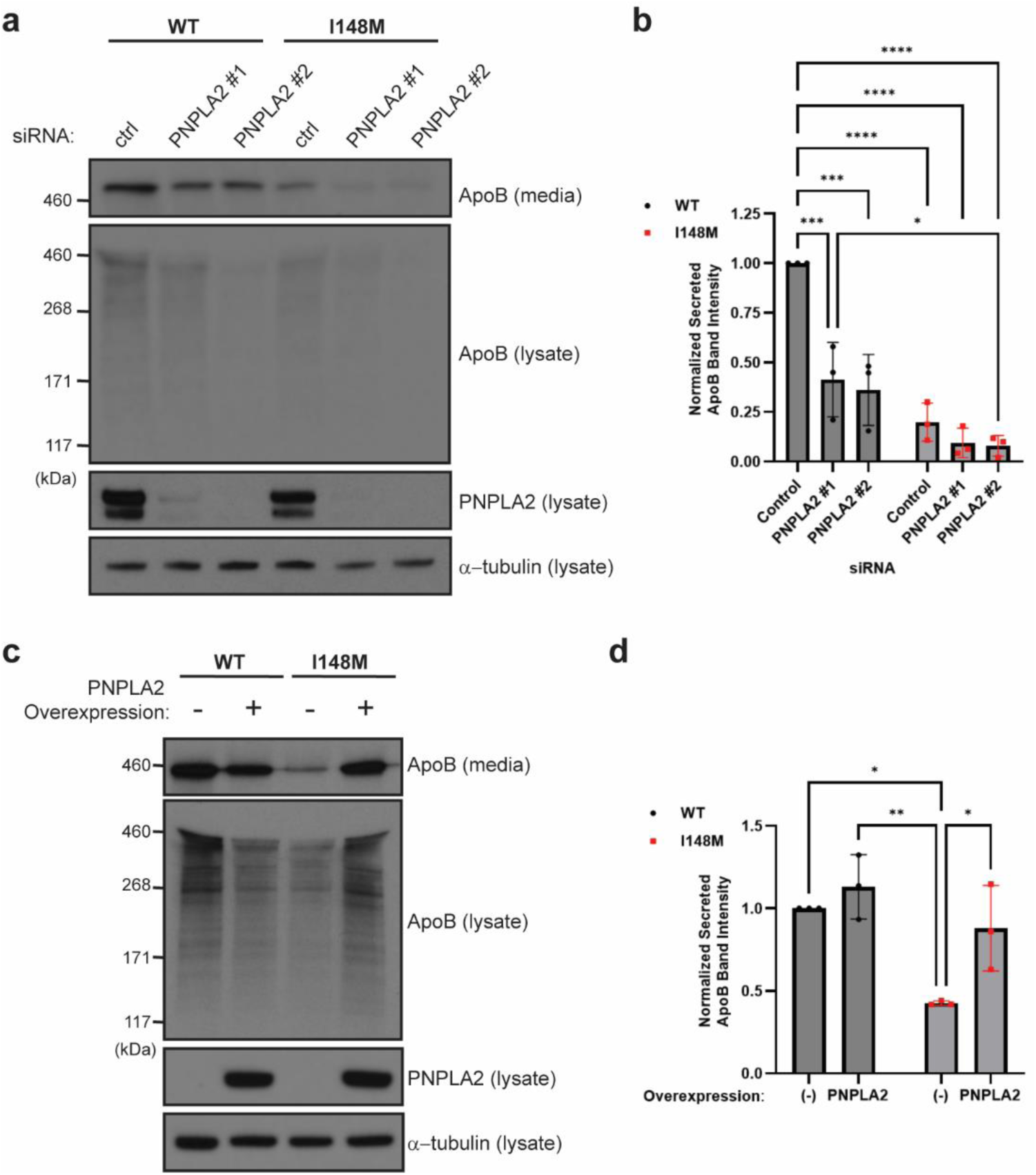
The ApoB secretory defect caused by PNPLA3-I148M is affected by PNPLA2 levels. **a,** Hep3B cells were treated for 72 h with control (non-targeting) or PNPLA2-targeting siRNA molecules. For the final 24 h, the media was exchanged for oleic acid (100 µM)-containing media. ApoB levels in the lysate and media were determined by immunoblots. **b,** ApoB immunoblots (**a**) were quantified by densitometry to determine relative ApoB levels in the media. Analyzed by two-way Anova with Tukey’s multiple comparisons test. WT control-I148M control, WT control-I148M PNPLA2 #1, and WT control-I148M PNPLA2 #2, adj. *p* < 0.0001; WT control-WT PNPLA2 #1, adj. *p* = 0.0006; WT control-WT PNPLA2 #2, adj. *p =* 0.0003; WT PNPLA2 #1-I148M PNPLA2 #2, *p* = 0.0435. *n* = 3 biological replicates. **c,** Parental Hep3B cells and Hep3B cells stably overexpressing PNPLA2 were treated as in **a**, and ApoB was detected in the lysate and media by immunoblotting. **d,** ApoB immunoblots (**c**) were quantified by densitometry to determine relative ApoB levels in the media. Analyzed by two-way Anova with Tukey’s multiple comparisons test. WT-I148M, adj. *p* = 0.0107; I148M-WT + PNPLA2, adj. *p* = 0.0031; I148M-I148M + PNPLA2, adj. *p* = 0.0367. *n =* 3 biological replicates. * *p* ≤ 0.05; ** *p* ≤ 0.01; *** *p* ≤ 0.001; **** *p* ≤ 0.0001.

**Figure 7:**
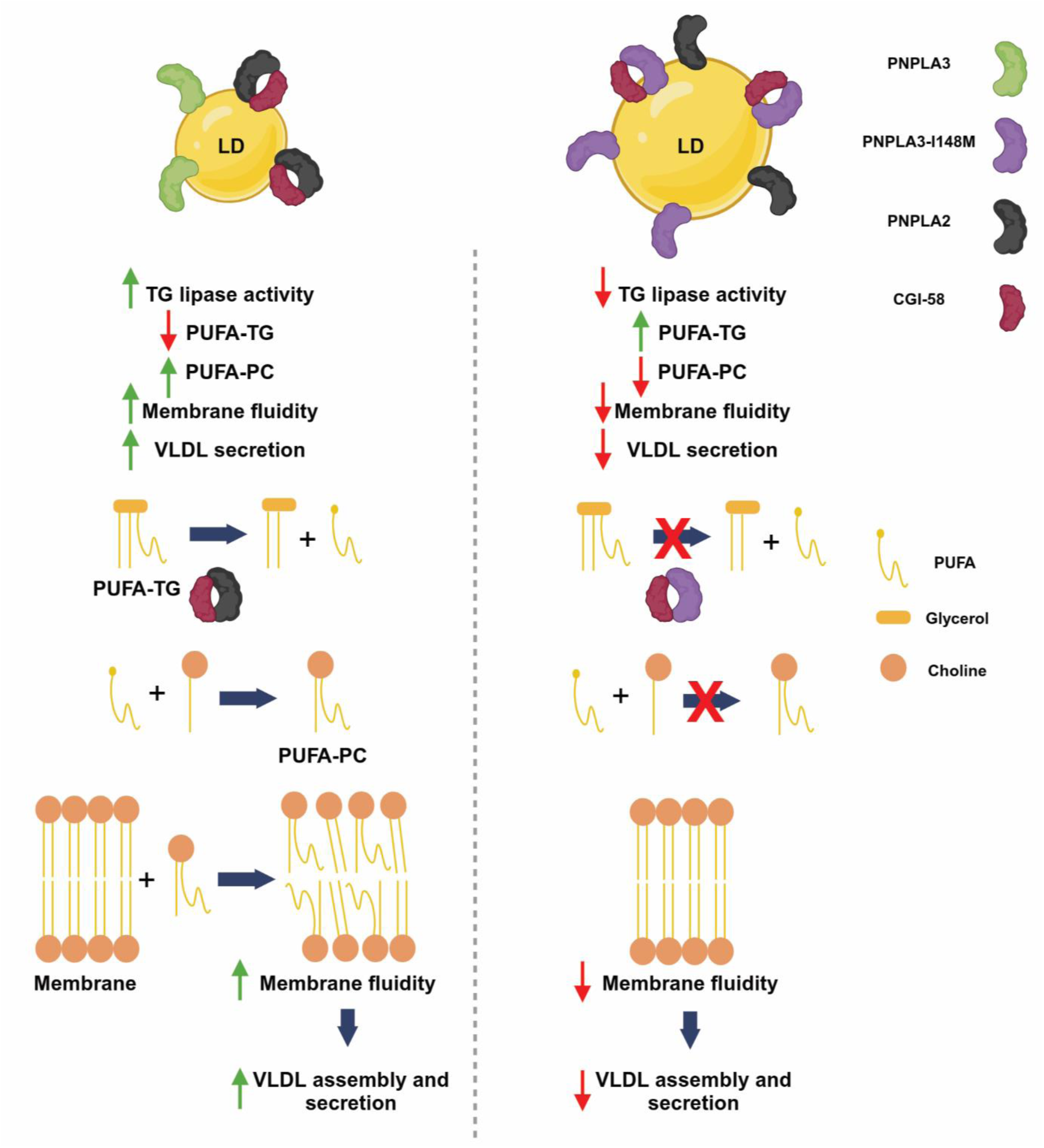
PNPLA3-I148M connects CGI-58-mediated lipolysis with VLDL biogenesis. Schematic diagram depicting the consequences of wild-type PNPLA3 expression (left) and PNPLA3-I148M expression (right). PNPLA3-I148M sequesters and inactivates CGI-58, impeding lipolysis by PNPLA2. Consequently, PUFAs remain sequestered in TG and are not available for incorporation into PC, reducing membrane dynamics and hindering VLDL biogenesis. In contrast, when WT PNPLA3 is present, homeostatic lipolysis occurs and allows for the proper balance of hepatic TG uptake/synthesis and breakdown/secretion.^†^

## Discussion

The defining characteristic of cells that express PNPLA3-I148M is that they accumulate aberrantly high levels of TGs stored in cytosolic LDs. The mechanism(s) by which PNPLA3-I148M causes such TG buildup, or steatosis, in humans, and downstream sequelae such as MASH and hepatocellular carcinoma, has remained elusive. Observations reported here and by others lead us to expand on an attractive model articulated by Hobbs, Cohen and colleagues. We propose a bipartite mechanism underlying the accumulation of LDs in I148M cells. As previously shown, and consistent with data reported here, PNPLA3-I148M sequesters CGI-58, an activator of the lipase activity of PNPLA2^15,16,27,64^. Both wild-type and I148M PNPLA3 bind CGI-58 and cells accumulate higher levels of the I148M variant, which may sequester CGI-58 with greater affinity^64^. In addition, PNPLA3-I148M•CGI-58 has diminished TG lipase activity^27,65,66^. Thus, I148M should sequester more CGI-58 than wild-type into a complex with less activity, resulting in less total (PNPLA2 + PNPLA3) lipase activity. This results in accumulation of TGs in the form of LDs. In addition, as shown in Fig. 4, I148M causes a shift in lipid species, with accumulation of PUFAs in TGs at the expense of the membrane phospholipid PC. This elicits a reduction in membrane dynamics, namely fluidity, which in turn leads to the second major consequence of I148M expression: altered formation and secretion of ApoB-containing VLDL particles. This manifests itself, depending on the experimental system, as reduced secretion of ApoB and/or secretion of VLDL particles of smaller size (i.e., fewer TGs) and altered lipid composition^18,31^. This leads us to propose that I148M causes steatosis by foreclosing two primary options the liver has for reducing excessive TG levels: metabolism by cytosolic lipase action^15,16,27,64^ and packaging into mature VLDL particles for secretion (Fig. 7). Consequently, TG-rich cytosolic LDs accumulate to very high levels.

There are at least two aspects of this model that still need to be reconciled. One is that the catalytically inactive S47A variant of PNPLA3 has been shown to mimic PNPLA3-I148M in mice, causing steatosis on a high-sucrose diet^66,67^. *Pnpla3* knockout mice do not resemble I148M in that they do not develop steatosis on a variety of diets, including a high-sucrose diet^20,21^. From these results, it appeared that altering the active site of PNPLA3 was not the same as losing expression of the protein. We found that, unlike I148M, PNPLA3-S47A does not affect ApoB secretion (Fig. 1c), and instead has normal ApoB secretion, similar to WT and PNPLA3 knockout cells. This discrepancy could be due to differences in the mouse and human proteins, or it could be due to structural or functional differences in the complexes that I148M and S47A PNPLA3 form with CGI-58. Future work is needed to dissect the differences between I148M and S47A, although this is not the first time that differences between the two variants have been observed. Mitsche *et al.* found that the hepatic LD lipidomic profile of I148M-knock-in mice differed from that of S47A-knock-in mice, the latter of which more closely resembled that of *Pnpla3*^-/-^ mice^68^.

A second aspect that needs to be reconciled is the role of WT PNPLA3 protein in LD metabolism and ApoB secretion under normal conditions. In the context of PNPLA2 knock-out mice on a lipogenic diet, PNPLA3 can serve as a triglyceride lipase that releases PUFAs from cytosolic LDs, allowing for their incorporation into membrane phospholipids for biogenesis of large VLDL particles^19^. While this may be the case under the specialized conditions employed in ref. 19, our data indicate that knockout or catalytic inactivation of PNPLA3 does not significantly impair ApoB secretion, suggesting that in the absence of I148M, PNPLA2 is the major source of PUFAs to sustain VLDL biogenesis. Indeed, depletion of PNPLA2 caused a 2.5-3-fold reduction in ApoB secretion (Fig. 6b). Uncertainty about the physiological function of the WT protein, coupled with observations that PNPLA3-I148M expression mimics PNPLA3 knockout in certain systems^18,69,70^, have led some to call PNPLA3-I148M a loss-of-function variant. However, this is inconsistent with the observation that PNPLA3 knockout mice do not develop steatosis^20,21^.

The relationship between PNPLA3 and VLDL secretion in mice and humans has been enigmatic for several years and riddled with conflicting results. Bulk TG secretion was not impaired over a two-hour period of lipoprotein lipase inhibition, following a four-to-six hour fast, in transgenic or knock-in I148M mice^22,65^. However, I148M knock-in mice on a corn-oil enriched Western diet exhibited impaired bulk TG secretion over a three-hour period of lipoprotein lipase inhibition^19^. A persistent challenge when comparing studies is that there are differences in expression of ApoB in humans and mice, as well as differences in sequence and expression of PNPLA3 itself.

The greatest challenge to the observations presented here come from human genetics studies reporting that I148M carrier status is not associated with altered total plasma TG or ApoB levels^1,2,17^. There are several issues that confound simple interpretation of these results. First, even between the human studies there are discrepancies. For example, radioisotope tracing studies with obese, non-diabetic men revealed reduced secretion of large VLDL particles in I148M carriers versus non-carriers^17^. Second, there are multiple sources of plasma TG and ApoB (e.g., chylomicrons secreted by the intestine), and human studies often fail to identify the specific form of ApoB detected (e.g., ApoB48 from the intestine vs. ApoB100 from the liver). Third, using the more sensitive technique of nuclear magnetic resonance to profile secreted lipoproteins, one group reported a striking reduction in large VLDL particles produced by insulin-resistant I148M carriers^31^. This same group reported changes in VLDL composition and the liver lipidome in homozygous I148M carriers similar to those reported here (e.g., increased TG-PUFA and decreased PC-PUFA in the liver) ^18^. Finally, one study with human homozygous carriers found similar levels of VLDL secretion, however this was in the face of a ∼ 3-fold increase in hepatic TG levels relative to non-carriers, suggesting that I148M carriers cannot effectively balance TG secretion with input levels^71^. Collectively, the human data combined with our results point to an inability of variant liver cells to metabolize TG and assemble and secrete VLDL particles at a rate that is sufficient to sustain appropriate TG homeostasis. Importantly, our findings establish, in controlled studies with isogenic cell lines, that altered VLDL biogenesis and ApoB secretion are cell-intrinsic consequences of the expression of endogenous I148M in cells of liver origin as opposed to other genetic, physiological, and/or environmental variables.

Future work will aim to address specifically where in the VLDL biogenesis process PNPLA3-I148M interferes. Following the generation of nascent VLDL particles in the ER, multiple studies have found evidence for generation in the Golgi of larger, mature particles that are closer in size and composition to those observed in circulation^37,72–74^. We previously found that PNPLA3-I148M associates with the Golgi and induces more LD-Golgi contacts^11^. A plausible model is that PNPLA3-I148M hinders formation and transfer of fatty acids at or proximal to the Golgi, disrupting the growth and trafficking of VLDL particles in the Golgi. LDs and the Golgi are rich in phospholipid remodeling enzymes that can generate PC-PUFA^75^. Therefore, PNPLA3-I148M would be well positioned to affect membrane dynamics, and consequently VLDL biogenesis, by inhibiting lipolysis at this node of the secretory pathway. We hypothesize that the variant can be limiting the supply of PUFAs for phospholipid remodeling enzymes via the regulation of lipolysis.

Given its importance in maintaining lipid homeostasis, lipolysis is a highly regulated process that involves multiple layers of regulation at the transcriptional and post-translational levels^76^. While the best characterized role for CGI-58 is in regulating PNPLA2 activity, several lines of evidence suggest that there could be PNPLA2-independent functions for CGI-58 in regulating lipolysis, including activation of WT PNPLA3. Consistent with this study, others have reported that CGI-58 overexpression alone, without co-expressing PNPLA2, reduces intracellular TG stores in cells and mice^15,30,61^. Modulating CGI-58 levels in cells has also been shown to impact VLDL biogenesis by regulating intracellular TG stores^61,62^, and CGI-58 knockdown can reduce VLDL-TG secretion in mice^63^. The role of PNPLA2 in VLDL biogenesis has been less clear than for CGI-58^77,78^. As the rate-limiting enzyme in TG lipolysis, PNPLA2 can catalyze the breakdown of TGs with a variety of fatty acyl chains, raising the question of why we observe, specifically, a decrease in PC-PUFAs in this study. We speculate that this relates to the fact that phospholipid remodeling enzymes have particular lipid specificities and show a preference for PUFA incorporation at the *sn-*2 position of phospholipids. Therefore, the outcome of a global deficiency in lipolysis is fewer substrates altogether for phospholipid remodeling, including PUFAs for the enzymes such as LPCAT3 that are needed to modulate membrane fluidity and dynamics^40,79^.

Acting at the intersection of two key pathways for TG breakdown and clearance – lipolysis and VLDL biogenesis – PNPLA3-I148M is uniquely positioned to cause the pleiotropic effects, ranging from LD accumulation to altered membrane dynamics and signaling, associated with MASLD. A phase 1 clinical trial of a PNPLA3-specific siRNA therapeutic to knockdown hepatic PNPLA3 levels showed promising results for reducing hepatic fat in homozygous carriers, representing a major step forward in the treatment of MASLD and further cementing the notion that the I148M variant is a neomorph as opposed to a loss-of-function^80^. However, given that we do not know the full spectrum of activities possessed by the WT and variant proteins, an allele-specific approach to targeting PNPLA3-I148M might be preferential^81^. Further investigation into role of PNPLA3-I148M and CGI-58 at the intersection of lipolysis and VLDL biogenesis, in addition to a more detailed molecular understanding of the complex formed by the two proteins, could open doors to novel therapeutic strategies for treating PNPLA3-I148M-driven MASLD and provide important insights into the mechanisms that regulate lipid metabolism in human health and disease.

## Methods

### Cell lines

The generation of paired Hep3B cells (PNPLA3 WT and I148M) by was previously described^11^. PNPLA3 knockout cells were generated by CRISPR gene editing of parental Hep3B cells at Synthego (Redwood City, CA, USA). PNPLA3-S47A cells were similarly generated from parental Hep3B WT and I148M cells by EditCo Bio, Inc. (Redwood City, CA, USA). Ribonucleoprotein complexes containing synthetic chemically-modified guide RNA were electroporated into the cells using EditCo’s optimized protocol. Editing efficiency was assessed upon recovery, 48 h post-electroporation. Genomic DNA was extracted from a portion of the cells, PCR-amplified and sequenced using Sanger sequencing. The resulting chromatograms were processed using EditCo’s Inference of CRISPR Edits (ICE) software (<u>ice.synthego.com</u>). To create monoclonal cell populations, edited cell pools were seeded at 1 cell/well using a single-cell printer into 96- or 384-well plates. All wells were imaged every 3 days to ensure expansion from a single-cell clone. Clonal populations were screened and identified using the PCR-Sanger-ICE genotyping strategy described above.

Hep3B cells were all cultured in Eagle’s Minimum Essential Medium (EMEM; ATCC) supplemented with 10% dialyzed fetal bovine serum (Gibco) and 1x Antibiotic-Antimycotic (Gibco). For siRNA treatment and lentiviral transduction, antibiotic was omitted from the media. CGI-58- and PNPLA2-overexpressing cell lines were generated by transducing Hep3B WT and I148M cells (250,000 cells/well of a 6-well plate) with lentivirus (Origene, CGI-58 RC201869L3V and PNPLA2 RC205708L3V), with a multiplicity of infection (MOI) of 10 and 10 µg/mL polybrene (Sigma, TR-1003-G). After 72 h, media was exchanged and cells were allowed to recover. The following day, media was supplemented with 2 µM puromycin dihydrochloride (Gibco, #A1113803) for selection. After 2 weeks of selection, cells were allowed to recover and propagate for use in experiments. Cells were split twice weekly to maintain at sub-confluence.

#### siRNA treatment

Hep3B cells were reverse-transfected in 6-well plates using Lipofectamine RNAiMAX Reagent (Invitrogen), according to the manufacturer’s instructions. 25 pmol of the following Silencer Select siRNA reagents (Thermo Scientific) were used per transfection: Negative Control siRNA #2 (4390846); CGI-58 s27425 and s224185; PNPLA2 s534784 and s534782. Cells were treated with siRNA for 72 h.

#### Immunoblotting

Western blots for proteins other than ApoB were performed as described^11^. For ApoB blots, clarified cell lysates and media (1:1 dilution in 2x SDS sample buffer with 5% β-mercaptoethanol) were loaded onto NuPAGE™ Tris-Acetate Mini Protein Gels, 3 to 8%, 1.0–1.5 mm (Invitrogen) and run in 1x NuPAGE™ Tris-Acetate SDS Running Buffer (Invitrogen) for 70 min at 150 V. The HiMark™ Pre-stained Protein Standard (Invitrogen, LC5699) was used because of the high molecular weight of ApoB. Gels were transferred onto 0.45 µm polyvinylidene difluoride membranes (ThermoFisher Scientific) for 16 h at 4°C (35 mA) using the Mini Trans-Blot Electrophoretic Transfer Cell (Bio-Rad). The remainder of the protocol was the same as for other protein blots^11^. All antibodies were diluted in phosphate-buffered saline (PBS) with 0.1% Tween-20. Primary antibodies used for immunoblotting: goat anti-apolipoprotein B (EMD, #178467; 1:500 dilution), BSA (Invitrogen, #A11133; 1:10,000 dilution), β-COP (Invitrogen, #PA1-061; 1:1,000 dilution); CGI-58 (Sigma, #WH0051099M1; 1:1,000 dilution); ε-COP (Sigma, #HPA043576; 1:1,000 dilution); FLAG (Sigma, F1804; 1:10,000 dilution); Nrf1 (Cell Signaling Technology, #8052; 1:1,000 dilution); PNPLA2 (Cell Signaling Technology, #2138); poly-ubiquitin (Enzo Life Sciences, #ADI-SPA-200-F; 1:10,000 dilution); Sec23A (Cell Signaling Technology, #8162; 1:1,000 dilution); Sec31A (Cell Signaling Technology, #13466; 1:1,000 dilution); α-tubulin (Cell Signaling Technology, #2144; 1:2,000 dilution). Secondary antibodies used for immunoblotting: anti-goat IgG-horseradish peroxidase (Bio-Rad, # 1721034); anti-mouse IgG-horseradish peroxidase (Bio-Rad, #1706516); anti-rabbit IgG-horseradish peroxidase (Bio-Rad, #1706515).

#### ApoB ELISA

Hep3B cells were treated with 100 µM oleic acid complexed with BSA (Sigma, O3008). After 24 h, media was removed and centrifuged, at 4°C, for 5 min at 500 x *g*. The supernatant was removed, supplemented with 100 µg/mL Pefabloc^®^ SC (Roche), and added to a 96-well ApoB ELISA plate (Invitrogen, EH34RB). Cells were either washed with Dulbecco’s Buffered Saline (DPBS; Gibco, #14190-136) and trypsinized for counting by Vi-Cell XR (Beckman Coulter) or washed with DPBS and lysed in RIPA buffer (Thermo Scientific, #89900) supplemented with cOmplete^™^ EDTA-free Protease Inhibitor Cocktail (Roche, #11836170001) and PhosSTOP^™^ Phosphatase Inhibitor (Roche, #4906845001). Lysate concentration was determined by *DC*^TM^ Protein Assay (Bio-Rad, #5000112), diluted 3-fold, and added to the 96-well ApoB ELISA plate. The ELISA was run according to the manufacturer’s instructions. Absorbance was determined using a SpectraMax® M5 Multi-Mode Microplate Reader (Molecular Devices) and SoftMax Pro Version 6.5.1 (Molecular Devices). Results were normalized either to cell count of concentration of lysate.

For Triacsin C treatment, 5 µM Triacsin C (Cayman Chemical, #10007448) dissolved in dimethyl sulfoxide (DMSO) was added to cell culture media for 24 h prior to oleic acid treatment (which also included 5 µM Triacsin C). For liposome treatment of cells, 100 µM liposomes made from PC(16:0/18:0) (Avanti Polar Lipids, #850456) or PC(16:0/20:4) (Avanti Polar Lipids, 850459) were added to media with 100 µM oleic acid-BSA for 24 h prior to performing the ELISA. Liposomes were generated and resuspended in DPBS as described^82^. For knockdown studies, media was exchanged for fresh media lacking antibiotics and supplemented with 100 µM oleic acid-BSA for 24 h prior to performing the ELISA. For CB-5083 treatment, media was supplemented with 5 µM CB-5083 (Cayman Chemical, #19311) in DMSO for the final 5 h of oleic acid treatment.

#### RNA analysis: qPCR

*APOB* expression determined by first isolating total RNA from Hep3B cells grown in 6-well plates using the RNeasy Mini Kit (Qiagen, #74104). Real-time qPCR was performed using the TaqMan^TM^ RNA-to-Ct^TM^ 1-Step Kit (ThermoFisher, 4392938) and the QuantStudio Real-Time PCR system (ThermoFisher, 7 Flex). The following primer-probe sets were used: Hs00181142_m1 (*APOB*) and Hs00427620_m1 (*TATA-box binding protein*, or *TBP,* used as a housekeeping control).

#### *Gaussia* luciferase assay

Hep3B cells were transfected with pCMV-*Gaussia*-Dura Luc (Thermo Scientific, #16191) for 24 h using Lipofectamine™ 3000 Transfection Reagent, according to the manufacturer’s protocol. Media was then replaced with fresh media with or without 5 µg/mL Brefeldin A (Sigma, B6542), in DMSO, for 24 h. Media was harvested and transferred to an opaque 96-well plate. The Pierce™ *Gaussia* Luciferase Glow Assay Kit (#16160) was used to measure luciferase activity with a SpectraMax® M5 Multi-Mode Microplate Reader, as described above. Cell viability in this study was determined by CellTiter-Glo® Luminescent Cell Viability Assay (Promega, G7573).

#### Lipidomics sample preparation

Hep3B WT and I148M cells were cultured in 6-well plates (1,000,000 cells/well) under basal conditions, or in media supplemented with 100 µM oleic acid complexed to BSA for 24 h. Cells were then washed in DPBS, trypsinized, washed again in DPBS, and snap frozen on dry ice. Lipids were extracted by resuspending the cell pellets in 100 µL water and 225 µL methanol, and the resulting suspension was transferred to a glass culture tube. 10 µL of lipidomics internal standard (SPLASH^®^ LIPIDOMIX^®^; Avanti Polar Lipids, #330707) was spiked into each sample, followed by vortexing and addition of 750 µL methyl-tert-butyl ether (MTBE). Samples were incubated on ice for 20 min, vortexed, and 188 µL water was added, followed by another round of vortexing. Samples were then centrifuged for 10 min at 3,200 x *g* and the top layer was transferred to a glass autosampler vial and evaporated under a stream of nitrogen gas. Dried samples were reconstituted in isopropanol and subjected to LC-MS for analysis. Three technical replicates were prepared for each sample, and there were three biological replicates.

#### Lipidomics LC-MS

5 µL of each lipid extract was subjected to LC-MS analysis using a Waters Acquity I-Class ultra-performance liquid chromatography system coupled to a Thermo Q Exactive HFX mass spectrometer. Lipids were separated on a Thermo Accucore C30 HPLC column (2.1 mm x 150 mm, 2.6 µm particle size). Mobile phase A (MPA) was 60:40 acetonitrile:water + 10 mM ammonium formate + 0.1% formic acid. Mobile phase B (MPB) was 90:8:2 isopropanol:acetonitrile:water + 10 mM ammonium formate + 0.1% formic acid. The initial mobile phase composition was 30% MPB and was increased to 43% MPB by 5 min, 50% MPB by 5.1 min, 70% MPB by 14 min, 99% MPB by 21 min, then held at 99% MPB until 24 min, followed by column equilibration at initial mobile phase conditions for 6 minutes. The mobile phase flow rate was maintained at 0.35 mL/min throughout the gradient and the column was maintained at 40°C. The mass spectrometer was operated in polarity switching mode to acquire data in both positive and negative mode from a single run. MS resolution was 60,000, AGC was 1e6, maximum IT was 200 ms, and the MS scan range was 200-1400 *m/z*. To annotate lipids, pooled QC samples were prepared with aliquots of each individual experimental sample and these were subjected to data-dependent MS2 analysis using the chromatography conditions detailed above. For data-dependent MS2 acquisition, the MS1 scan was set to 30,000 resolution, 1e5 AGC, 50 ms maximum IT, isolation window of 1 *m/z*, collision energy of 30, and dynamic exclusion of 10 s. Annotated lipids were incorporated into a compound database, and this database was used to integrate peak features in the MS1 sample data using TraceFinder Software (Thermo Fisher Scientific).

#### Phosphatidylcholine quantification

Hep3B cells (WT and I148M) were grown in 6-well plates. Half of the wells were treated with 100 µM oleic acid conjugated to BSA for 24 h. All cells were washed three times with cold DPBS prior to lysis. Cells were lysed and PC content was quantified using the Phosphatidylcholine Assay Kit (Abcam, ab83377), according to the manufacturer’s instructions.

#### Fluorescence microscopy

For LD imaging with Triacsin C treatment, cells were seeded on 8-well Nunc Lab-Tek chambered coverglass (ThermoScientific, #155409). Following 24 h treatment with Triacscin C, media was removed and cells were washed with PBS, fixed, and stained with 2 µM Hoechst 33342 (ThermoScientific, # 62249) and HCS LipidTOX Green Neutral Lipid Stain (Invitrogen, # H34475) in PBS for 30 min at room temperature. Cells were washed and imaged using an inverted Zeiss LSM 800 confocal microscope and ZEN 2.1 software. Confocal images were obtained using an oil immersion Plan-Apochromat 63x/1.4 NA DIC objective lens.

For LD quantification during starvation, Hep3B cells were seeded into 96-well black CellCarrier tissue culture plates with optically clear bottoms (PerkinElmer, #6005550). Cells were incubated overnight so that they can settle, then media was exchanged to media supplemented with 200 µM oleic acid complexed with BSA. After 18 h, cells were washed twice with DPBS and stained with 2 µM Hoechst 33342 and HCS LipidTOX Green Neutral Lipid Stain in PBS for 30 min at room temperature. Then, staining solution was removed and replaced with Earle’s Balanced Salt Solution (EBSS; Gibco, #14155063), with or without 100 nM Bafilomycin A1 (Sigma, #19-148) in DMSO. Live cell imaging was conducted using the Operetta CLS high-content imaging system (PerkinElmer). Cells were kept at 37°C in a humidified, 5% CO_2_ compartment during imaging, and images were acquired at 0, 2, and 4 h of EBSS incubation using a 63x water immersion objective. Cells were segmented and LDs were quantified using Harmony 4.9 software.

#### Generalized polarization analysis

For generalized polarization studies, Hep3B cells were seeded into 8-well Nunc Lab-Tek chambered coverglass, as above, and incubated overnight. The following day, media was exchanged for Opti-MEM media (Gibco) with 5 µM di-4-ANEPPDHQ (Invitrogen, #D36802) in DMSO. Cells were incubated at 37°C for 30 min, then imaged at 37°C on an SP8 confocal microscope (Leica) using a 63x/1.3 NA glycerol immersion lens. Image acquisition parameters, including for calibration images, were as described in Owen *et al*^83^. Briefly, di-4-ANEPPDHQ fluorescence was excited using the 488 nm laser line and the emission was captured simultaneously in the ranges of 500-580 nm and 620-750 nm for the two channels.

The analysis of generalized polarization images was carried out using the Generalized Polarization Plugin in ImageJ 1.54g^84^. GP values were calculated using the formula GP = (*I*_500-580_ – G\**I*_620-750_)/(*I*_500-580_ + G\**I*_620-750_), where *I* represents the intensity in each pixel in the image acquired in the indicated channel and G is the calibration factor. Since GP values obtained from the fluorescence images strongly depend on instrumental factors, such as filter settings and gain, the GP values were calibrated with the calibration factor G. The G-factor was calculated by correcting the GP value from the microscope images of a 5 µM di-4-ANEPPDHQ solution by the GP value obtained by fluorometric measurements of the solution using a SpectraMax® M5 Multi-Mode Microplate Reader at appropriate emission wavelengths (excitation at 488 nm, emission at both 560 nm and 650 nm).

#### Label-free live cell holotomographic imaging

Live-cell, label-free LD imaging in Hep3B cells overexpressing CGI-58-FLAG was done using the Nanolive CX-A 3D Cell Explorer microscope (Nanolive), as previously described^11^. Briefly, parental Hep3B cells and Hep3B cells stably overexpressing CGI-58-FLAG were seeded onto 96-well glass-bottom plates in phenol red-free media and treated with 200 µM oleic acid. After 1 h of plate thermalization, refractive index images were acquired for 16 h at a 10 min interval using a 60x/0.8 NA objective and 3 x 3 GridScan mode. Cells and LDs in refractive images were segmented and quantified using the Smart Lipid Droplet Assay module (Version 2.0) in the artificial intelligence/machine learning-powered Eve Analytics software (version 2.1.0.1987).

### Primary human hepatocytes

Primary hepatocytes were obtained from Lonza (lots #4056B and 4111C) or BioIVT (lots BVI, BXM, FZQ, IXL, and ONR). PNPLA3 variant status in lots was confirmed by an allele-specific TaqMan assay, as described^11^. Hepatocytes were plated in hepatocyte plating media (HPM): Dulbecco’s Modified Eagle’s Medium (DMEM; Gibco) with 2 mM L-glutamine (Gibco), 1x insulin-transferrin-selenium (ITS; Gibco, #41400045), and 10% fetal bovine serum (Gibco). Hepatocytes were maintained in hepatocyte incubation media (HIM): William’s Medium E (Sigma, #W1878), 2 mM L-glutamine, 1x ITS, and 50 U/mL penicillin-streptomycin (Gibco, #15070063). Frozen vials (1 mL) of primary hepatocytes were quickly thawed (< 90 s) in a 37°C water bath and diluted into 5 mL pre-warmed HPM. Cells were mixed by gently inverting the tube 3 times. Cells were counted on a microscope with 10X magnification using a hemocytometer after diluting a small sample in 0.4% Trypan Blue (Gibco) to determine viability. Cells were diluted in pre-warmed HPM to a concentration of 700,000 viable cells/mL, and 1 mL of cell suspension was added to each well of a 12-well Collagen I-coated plate (STEMCELL Technologies, #100-0363). Cells were allowed to settle overnight in a humidified incubator at 37°C with 5% CO_2_.

#### ApoB ELISA

After plating and incubation overnight, media was exchanged for 0.6 mL HPM supplemented with or without 200 µM oleic acid complexed with BSA and incubated for 48 h. Media was removed and centrifuged at 300 x *g*, for 5 min at 4°C, and then the supernatant was supplemented with 1x Halt™ Protease-Phosphatase Inhibitor cocktail (ThermoFisher, #78440) for analysis by ApoB ELISA (Invitrogen, EH34RB). Cells in 12-well plate were washed twice with cold DPBS supplemented with 1x protease-phosphatase inhibitor cocktail and then lysed directly in 50 µL RIPA buffer supplemented with Halt™ Protease Inhibitor Cocktail (Thermo Scientific, #87785). Lysate concentration was determined by *DC*^TM^ Protein Assay. Lysates were then diluted 6x and used for the ApoB ELISA. The ELISA was performed as described above for Hep3B cells.

For *PNPLA3* knockdown studies, two WT lots (4056B and BVI) and two I148M lots (FZQ and ONR) of primary hepatocytes were plated on 24-well Collagen I-coated plates (Gibco, # A1142802), and media was exchanged to HPM (500 µL/well) 4 h later instead of after an overnight incubation. Media was then exchanged the next day to 250 µL fresh HIM, and cells were transfected with 1 nM control siRNA (ON-TARGETplus Non-Targeting Control siRNA #1; Horizon, #D-001810-01-05) or *PNPLA3* siRNA (made by Amgen; sense strand: CAACGUACCCUUCAUUGAUU, antisense strand: AUCAAUGAAGGGUACGUUGUU) using Lipofectamine RNAiMAX Reagent, according to the manufacturer’s instructions. After 48 h, media and cell pellets were collected and prepared for ApoB ELISA or for dPCR, as previously described^11^.

### Animals

All animal procedures described herein were approved by the Institutional Animal Care and Use Committee (IACUC) of Charles River Accelerator and Development Lab (CRADL, AAALAC accredited) and cared for in accordance with the *Guide for the Care and Use of Laboratory Animals*, 8th Edition (National Research Council (U.S.)). Mice were group-housed in a climate-controlled room at 22±2°C with a twelve-hour light (0600-1800), followed by twelve-hour darkness cycle. Animals had *ad libitum* access to a regular chow diet (Inotiv, 2920X) and were caged in irradiated Innovive Disposable IVC Rodent Caging Systems (San Diego, CA) with pre-filled sterile and acidified water bottles. As indicated in the text, for some experimental conditions, mice had *ad libitum* access to the ALIOS diet during study (Teklad, TD.190883).

Transgenic human PNPLA3-I148M knock-in (huPNPLA3-I148M KI) mice were developed by Inotiv (Maryland Heights, MO) for Amgen and the colony was both maintained and genotyped at Charles River Laboratories (CRL, Hollister, CA). The huPNPLA3-I148M KI mouse line was created using gene-editing technology to remove the wild-type (WT) C57BL/6J mouse *Pnpla3* gene and insert the full-length coding region of the huPNPLA3-I148M gene. A phenotypic characterization study was performed comparing WT and homozygous huPNPLA3-I148M littermates; results of this characterization study indicated no significant genotype-related phenotypic differences between the two sets of littermates.

Eighteen-to-twenty-week-old male huPNPLA3-I148M KI homozygous (huI148M/huI148M; HOM) mice, wild-type mice (msI148I/msI148I; WT), and heterozygous mice (huI148M/msI148I; HET) were allowed to acclimate at Amgen four weeks prior to harvest.

For all *in vivo* studies described, mice were fasted overnight prior to study termination. Before harvest, mice were weighed and then anesthetized with isoflurane (NDC, 66794-0017-25). From each animal, blood was collected via intracardiac puncture using a 1 mL 25-gauge Tuberculin syringe and transferred to EDTA-coated microtainer tubes (BD, #365974), inverted eight-to-ten times, and stored on ice for no more than 30 min. Mice, still under deep anesthesia, were then euthanized by a secondary physical method and the left and right liver lobes were collected, weighed, and immediately snap frozen in liquid nitrogen until processing and analysis. Filled EDTA-coated tubes were centrifuged, at 4°C, for 15 min at 3500 rpm in a microcentrifuge, and then transferred to new EDTA-microtainer tubes, which were centrifuged at 4000 rpm for 15 min. Plasma was then transferred to Protein LoBind® tubes (Eppendorf, #022431081). A 100X stock Halt™ Protease-Phosphatase Inhibitor Cocktail (ThermoFisher, #78440) was added (1% v/v) to each sample. Each tube was vortexed briefly and centrifuged at low speed to collect plasma at the bottom of the tube, and then samples were transferred to a 96-well plate (Corning, #3752), sealed, frozen, and stored at -80°C.

#### RNA analysis: dPCR and qPCR

RNA was extracted from snap-frozen liver by homogenizing in TRIzol™ Reagent (ThermoFisher, #15596026), followed by chloroform extraction. Lysate samples were centrifuged for 15 min at 12,000 × *g*, at 4°C, and the aqueous phase containing the RNA was transferred to a new tube for RNA isolation using the QIAcube System (Qiagen, #9001292) and RNeasy Mini Kit reagents (Qiagen, #74106), both according to the manufacturer’s instructions. Isolated RNA samples were analyzed using a QIAxpert system. Samples were subsequently treated with RQ1 RNase-Free DNase (Promega, #M6101) and prepared for digital PCR using the High-Capacity cDNA Reverse Transcription Kit (Applied Biosystems, #4368813) or real-time qPCR using the TaqMan^TM^ RNA-to-Ct^TM^ 1-Step Kit (ThermoFisher, #4392656). Digital PCR was performed using a C1000 Touch™ Thermal Cycler (Bio-Rad, #1851197) and Real-time qPCR was performed using a QuantStudio Real-Time PCR system (ThermoFisher, 7 Flex). Results are presented as the relative fold change in mRNA expression, normalized to the housekeeping gene, mouse *TATA-box binding protein* (m*Tbp*). All samples were run in duplicate.

Primer-probe sets used for dPCR: huPNPLA3 (Invitrogen, Hs00228747_m1); msPnpla3 (Invitrogen, Mm00504420); Tbp (IDT, Mm.PT.39a.22214839).

Primer-probe sets used for qPCR: Acca (Invitrogen, Mm01304257_m1); ApoB (Invitrogen, Mm01545150_m1); Fdft1 (Invitrogen, Mm01598574_g1); Hmgcr (Invitrogen, Mm01282499_m1); Mia2 (Invitrogen, Mm00616368_m1); Mttp (Invitrogen, Mm00435015_m1); Pnpla2 (Invitrogen, Mm00503040_m1); Pnpla3 (Invitrogen, Mm00504420_m1); PNPLA3 (Invitrogen, Hs00228747_m1); Scd1 (Invitrogen, Mm00772290_m1); Srebf1 (Invitrogen, Mm00550338_m1); Tmem41b (Invitrogen, Mm00781053_s1); Tbp (Invitrogen, Mm01277042).

#### Triglyceride and cholesterol content

To determine hepatic triglyceride and cholesterol content, approximately 50 mg of frozen liver tissue from each animal was weighed on ice and then homogenized independently in a 2 mL Eppendorf™ Snap-Cap Microcentrifuge Safe-Lock™ Tube (ThermoFisher, #05-402-8) with a 5mm stainless steel bead (Qiagen, #69989) and 1 mL of isopropyl alcohol (Sigma, #I9526). Samples were lysed using a TissueLyser II (Qiagen, #85300) for 2 min and then incubated on ice for approximately 1 h, with 2 to 3 rounds of vortex during the incubation. After cold incubation, samples were centrifuged at 10,000 rpm x 15 min at 4°C, and then the supernatant was transferred to a new tube. Triglyceride concentration was determined using the Infinity^TM^ Triglyceride reagent (ThermoFisher, #TR22421) and Triglyceride standard (Pointe Scientific, #T7531-STD) according to the manufacturer’s instructions. Cholesterol concentration was determined using the Infinity^TM^ Cholesterol reagent (ThermoFisher, #TR13421) and Cholesterol standard (ThermoFisher, #23-666-198) according to the manufacturer’s instructions. Absorbance was determined using a SpectraMax® Plus 384 Microplate Reader (Molecular Devices) and SoftMax Pro Software, Version 6.5.1 (Molecular Devices).

#### Lipoprotein isolation by size-exclusion chromatography

Plasma lipoprotein particles were isolated using an AKTA Purifier FPLC System. Sample volumes from 100 – 450 µL were loaded onto a Superose 6 Increase 10/300 GL column (Cytiva, #29-0915-96) using an A-905 Autosampler (Amersham).

Fractionation was performed in PBS with 3 mM EDTA using a flow rate of 0.5 mL/min. 200 – 400 µL fractions were collected in 96-well plates using a Frac-950 automated fraction collector (GE AKTA). Additional separation resolution on large particles was accomplished utilizing the same protocol with a Sephacryl S-500 HR column (Cytiva, #17-0613-10). Fractions were subject to analytic assays to measure cholesterol, TG, and ApoB content.

#### ApoB ELISA

Plasma apolipoprotein B (ApoB) was determined using a mouse ApoB ELISA kit (Abcam, ab230932) according to the manufacturer’s instructions. Absorbance was determined using a SpectraMax® Plus 384 Microplate Reader, as described above.

### Data presentation and statistical analyses

All data were graphed and analyzed in GraphPad Prism 10 (Dotmatics). For ELISA data below the detectable range of the ApoB standard curve, values were considered “0.” Structural depictions were generated in PyMOL Version 2.5.5 (Schrödinger).

## Author Contributions

D.J.S., I.C.R., and R.J.D. conceived of the studies and analyses for the manuscript. D.J.S. designed, performed, and analyzed cellular studies. L.L. conducted and analyzed qPCR/dPCR experiments and ApoB secretion studies with primary human hepatocytes. E.L.L.G. conducted and analyzed lipidomics studies. P.F. performed and analyzed lipoprotein fractionation from mouse plasma. D.J.S., J.X., and J.F. designed and evaluated microscopy experiments and platforms. D.J.S. and J.X. performed and analyzed microscopy studies. I.C.R. and D.J.S. designed *in* vivo studies. D.P. and J.B. conducted the *in vivo* studies and performed *ex vivo* analysis. I.C.R. and R.J.D. coordinated and oversaw research and manuscript preparation. D.J.S. wrote the manuscript with input from co-authors.

## Acknowledgements

We thank the Amgen Postdoc Program and Amgen Research for funding and support for this work. We thank Michelle Chen and Devin Wakefield for Operetta data analysis support. We thank Justin Murray and Bryan Meade for providing the *PNPLA3* siRNA. We thank Rati Verma, Simon Jackson, Saptarsi Haldar, and Stephen Lee for helpful scientific discussions and support while conducting the experiments for and preparing this manuscript. We thank Jennifer Mamrosh and members of the Deshaies Lab for input and feedback during the early stages of this project.

## Footnotes

† Created in BioRender. **Fig. 1g**: Sherman, D. (2024) BioRender.com/e09p999; **Fig. 2a, f, and Fig. S5a:** Sherman, D. (2024) BioRender.com/t23s872; **Fig. 3a**: Sherman, D. (2024) BioRender.com/c40i553; **Fig. 7**: Sherman, D. (2024) BioRender.com/r41a327.

