## Supplementary Data for "PNPLA3-I148M is a Neomorph that Interferes with Two Primary Hepatic Triglyceride Clearance Pathways"

[1] Amgen Research, Thousand Oaks, CA 91320; [2] Amgen Research, South San Francisco, CA 94080; [3] Co-senior authors

**This file includes:**

Supplementary Figures 1-6
Supplementary Table 1
Footnote

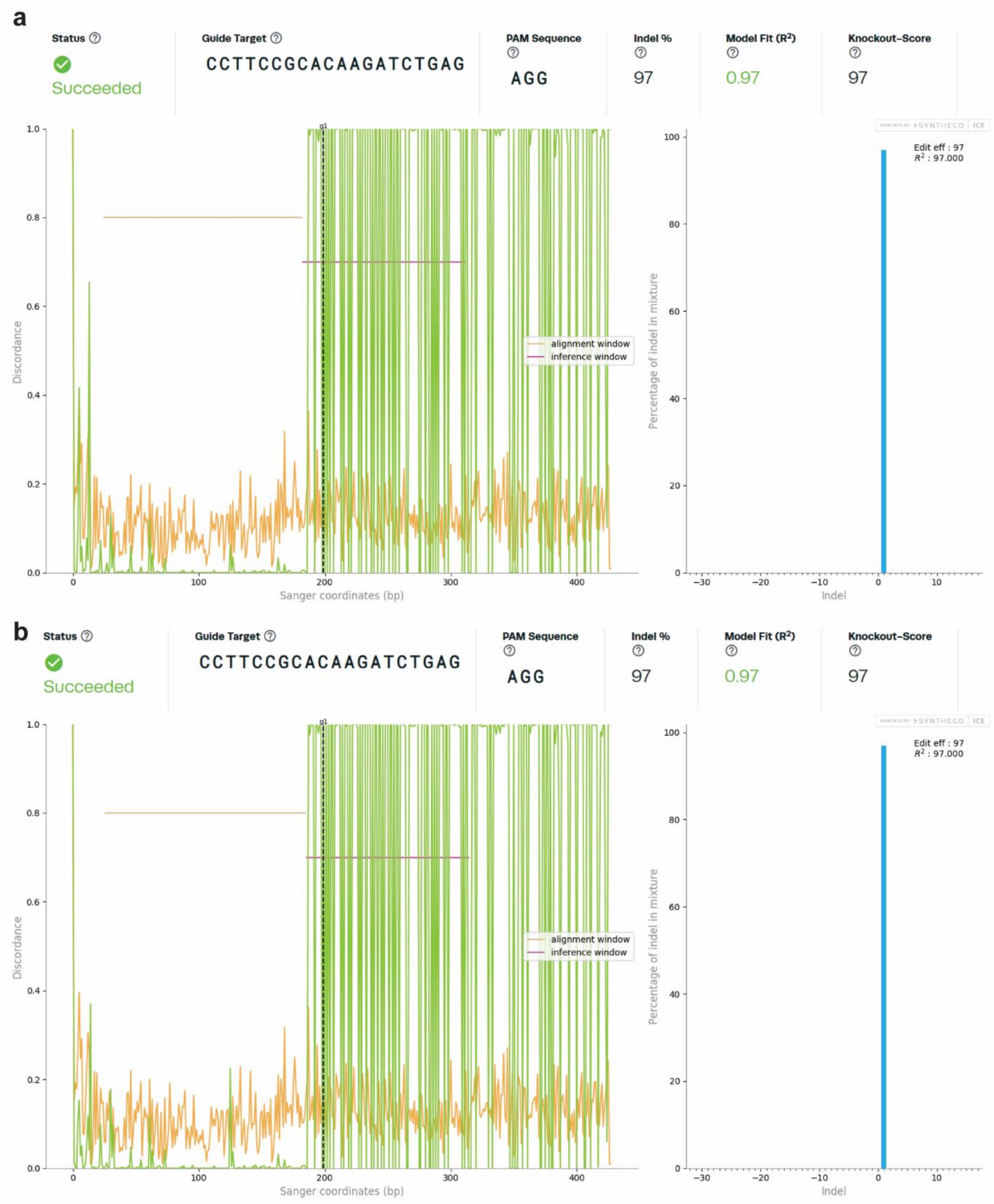


**Figure S1: EditCo's Inference of CRISPR Edits (ICE) Analysis of Hep3B *PNPLA3* Knockout cells.** Table of guide target, PAM sequence, % indels, model fit ($R^2$), and knockout scores for two clonal populations (**a,b**) of Hep3B *PNPLA3* knockout cells. Model fit ($R^2$) is how well the Sanger sequencing results of the edited clone fit the predicted indel distribution. Knockout score is the proportion of indels that indicate a frameshift, or those that are 21+ base pairs in length. Below the tables are graphs that show the level of disagreement between the trace file of the edited sample relative to the control sequence (i.e., discordance; left) and the percentage of indels in the edited mixture (right).

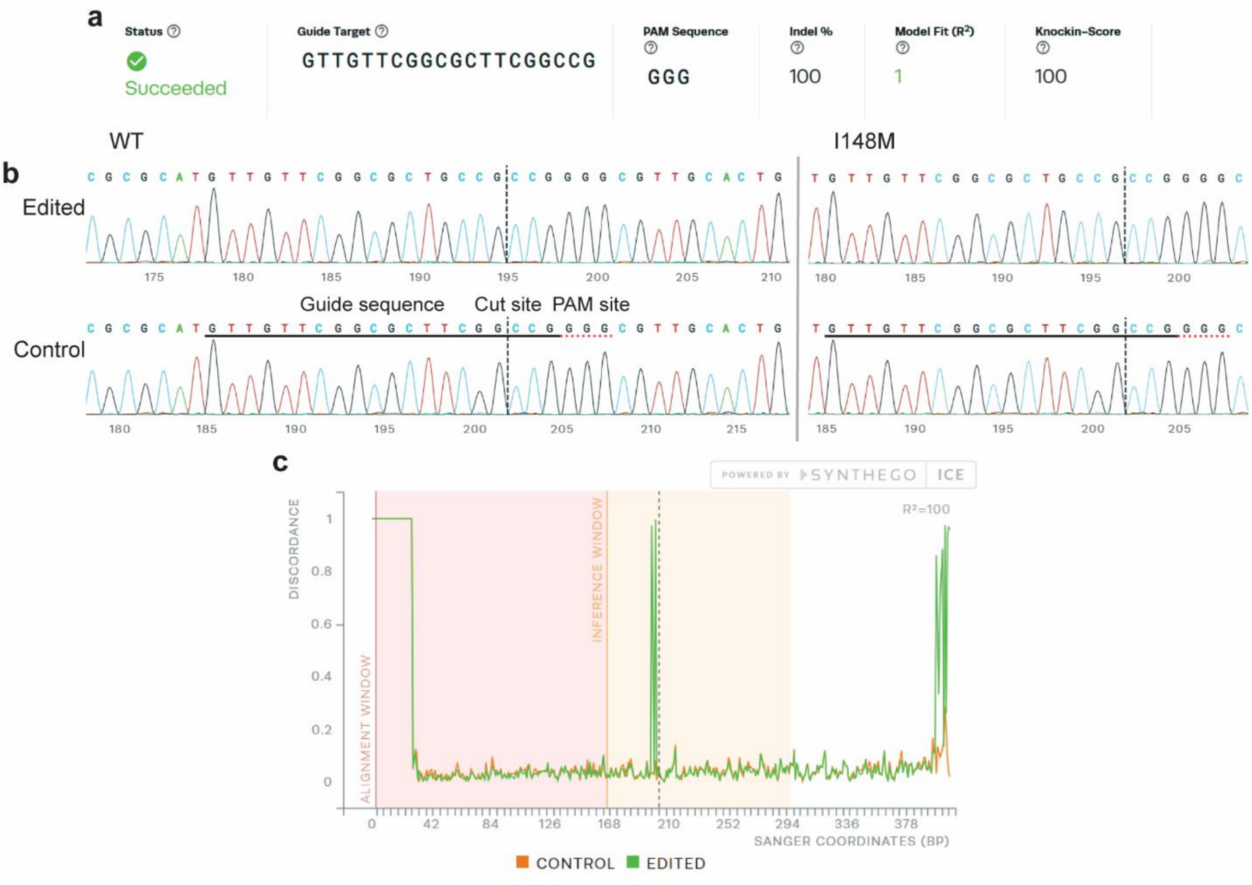


**Figure S2: EditCo's Inference of CRISPR Edits (ICE) Analysis of Hep3B PNPLA3-S47A cells.**
**a,** Table of guide target, PAM sequence, % indels, model fit ($R^2$), and knock-in score for Hep3B PNPLA3-S47A knock-in cells. Model fit ($R^2$) is how well the Sanger sequencing results of the edited clone fit the predicted indel distribution. Knock-in score is the proportion of indels that indicate a knock-in insert. Results were the same for gene-edited WT and I148M Hep3B cells. **b,** Sanger sequencing trace results for the region surrounding the S47A edit in WT Hep3B (left) and I148M Hep3B (right) cells. Guide sequence, cut site, and PAM site are labeled in the traces. **c,** Discordance plot of Hep3B PNPLA3-S47A cells shows the level of alignment per base between control and the edited samples in the inference window (the region around the cute site). The green and orange lines should be close together before the cut site, with editing resulting in discordance. Similar results were obtained for I148M cells.

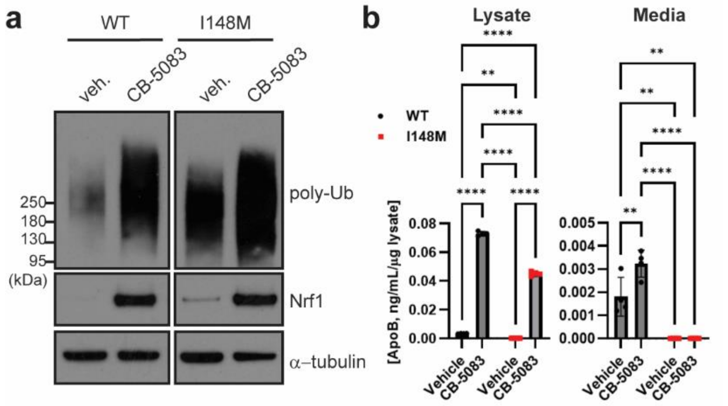


**Figure S3: PNPLA3-I148M causes degradation of intracellular ApoB in Hep3B cells. a,** Representative poly-ubiquitin and NFE2L1/Nrf1 immunoblots of vehicle- or CB-5083-treated Hep3B cells (WT and I148M). **b,** ELISA quantification of ApoB in the lysate and media of oleic acid (100 µM)-treated Hep3B cells. Cells were treated with vehicle (DMSO) or 5 µM CB-5083 for the final 5 h of oleate treatment. Analyzed by two-way Anova with Tukey's multiple comparisons test. Lysate: WT vehicle-I148M vehicle, adj. $p = 0.0034$; all other comparisons, adj. $p < 0.0001$. Media: WT vehicle-WT CB-5083, adj. $p = 0.0088$; WT vehicle-I148M vehicle, adj. $p = 0.0014$; WT vehicle-I14M CB-5083, adj. $p = 0.0014$; I148M vehicle-WT CB-5083 and WT CB-5083-I148M CB-5083, adj. $p < 0.0001$. $n = 3$ biological replicates. ** $p \leq 0.01$; **** $p \leq 0.0001$.

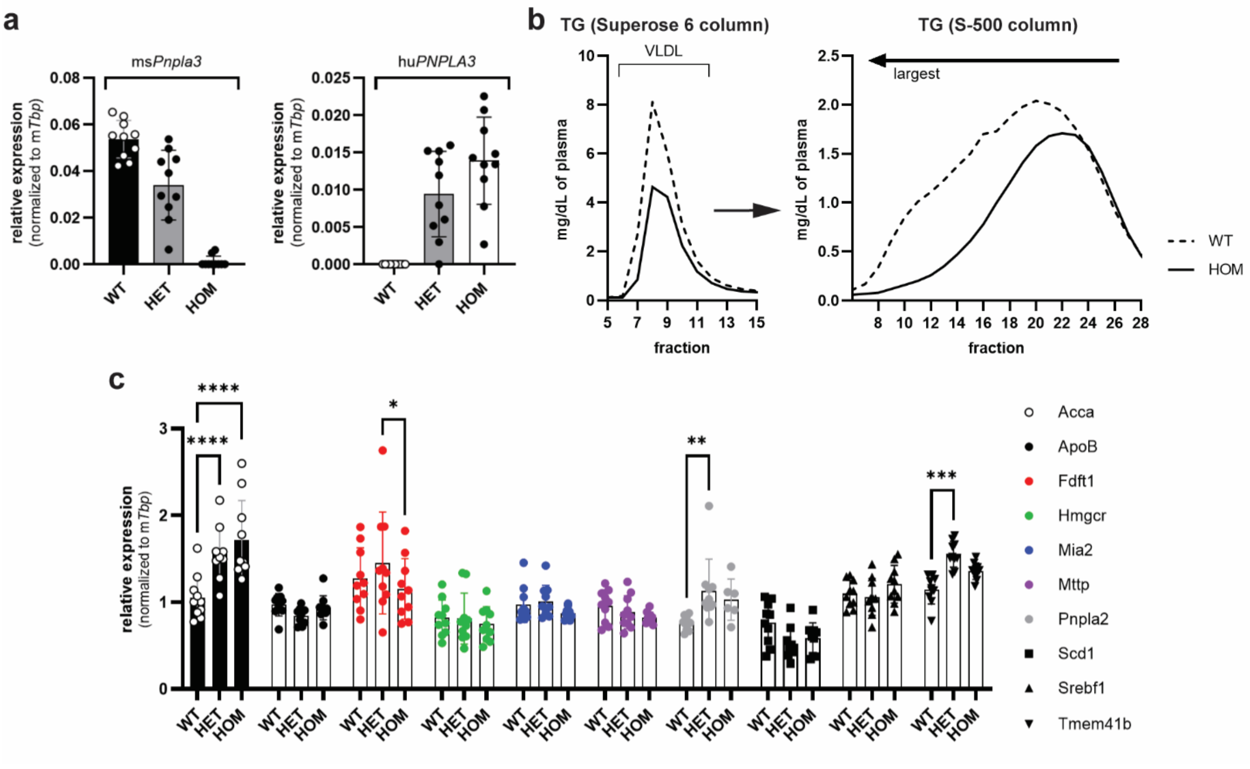


**Figure S4: Characterization of liver gene expression and VLDL-TG secretion in PNPLA3 mouse models. a,** dPCR analysis of mouse *Pnpla3* and human *PNPLA3* expression in the livers of endogenous knock-in mouse models. Expression levels were normalized to the mouse housekeeping gene *Tbp*. **b,** Following an overnight fast, plasma from 6 wild-type (WT) and 25 homozygous huPNPLA3-I148M knock-in (HOM) mice on a CHOW diet was collected, pooled, and subjected to size-exclusion chromatography on a Superose 6 column (left) and a Sephacryl S-500 column (right). TG in each fraction was quantified. **c,** qPCR analysis of liver lysates to determine expression of various genes involved in lipid metabolism and VLDL biogenesis. Analyzed by two-way Anova with Tukey's multiple comparisons test. *Acca* WT-HET and WT-HOM, adj. $p < 0.0001$; *Fdft1* HET-HOM, adj. $p = 0.0191$; *Pnpla2* WT-HET, adj. $p = 0.0026$; *Tmem41b* WT-HET, adj. $p = 0.0007$. * $p \leq 0.05$; ** $p \leq 0.01$; *** $p \leq 0.001$; **** $p \leq 0.0001$.

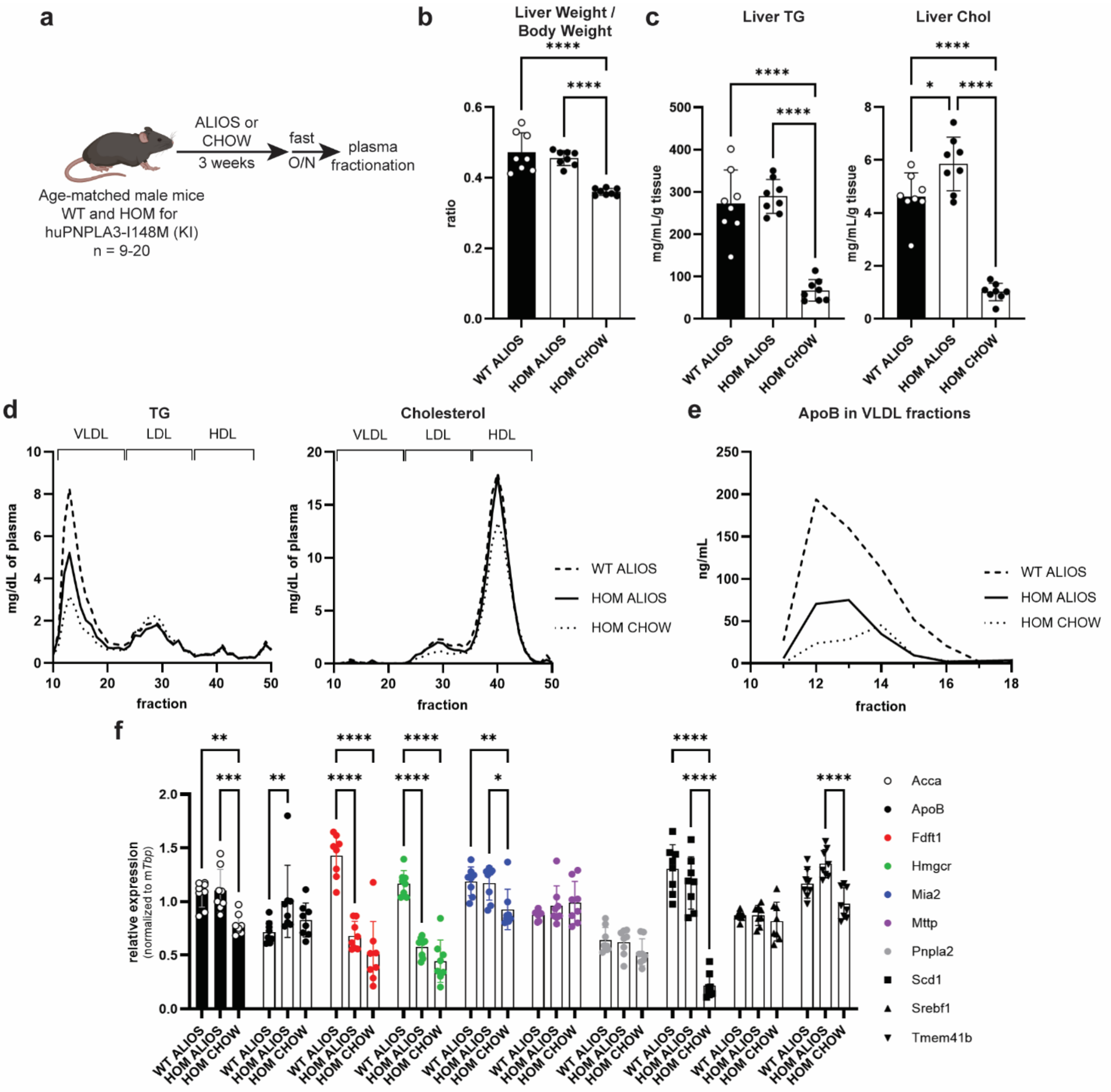


**Figure S5: huPNPLA3-I148M knock-in mice have a VLDL secretory defect on the ALIOS diet. a,** Schematic diagram of homozygous huPNPLA3-I148M knock-in (HOM) mouse study on the ALIOS diet.† **b,** Liver weight-to-body weight ratio of mice. Analyzed by ordinary one-way Anova with Tukey's multiple comparisons test. WT ALIOS-HOM CHOW and HOM ALIOS-HOM-CHOW, adj. $p < 0.0001$. **c,** Quantification of liver triglyceride and liver cholesterol content. Analyzed by ordinary one-way Anova with Tukey's multiple comparisons test. Liver TG: WT ALIOS-HOM CHOW and HOM ALIOS-HOM-CHOW, adj. $p < 0.0001$. Liver chol: WT ALIOS-HOM ALIOS, adj. $p = 0.0156$; WT ALIOS-HOM CHOW and HOM ALIOS-HOM-CHOW, adj. $p < 0.0001$. **d,** Pooled mouse plasma following an overnight fast was subject to size-exclusion chromatography on a Superose 6 column and TG and Chol levels were quantified in the fractions. **e,** VLDL fractions were analyzed by ApoB ELISA. **f,** qPCR analysis of liver lysates to determine expression of various genes involved in lipid metabolism and VLDL biogenesis. Analyzed by two-way Anova with Tukey's multiple comparisons test. *Acca* WT ALIOS-HOM CHOW, adj. $p < 0.0023$; *Acca* HOM ALIOS-HOM CHOW, adj. $p = 0.0006$; *ApoB* WT ALIOS-HOM ALIOS, $p = 0.0021$; *Fdft1* WT ALIOS-HOM ALIOS and WT ALIOS-HOM CHOW, adj. $p < 0.0001$; *Hmgcr* WT ALIOS-HOM ALIOS and WT ALIOS-HOM CHOW, adj. $p < 0.0001$; *Mia2* WT ALIOS-HOM CHOW, adj. $p = 0.0057$; *Mia2* HOM ALIOS-HOM CHOW, adj. $p = 0.0109$; *Scd1* WT ALIOS-HOM CHOW and HOM ALIOS-HOM CHOW, adj. $p < 0.0001$; *Tmem41b* HOM ALIOS-HOM CHOW, adj. $p < 0.0001$. * $p \leq 0.05$; ** $p \leq 0.01$; *** $p \leq 0.001$; **** $p \leq 0.0001$.

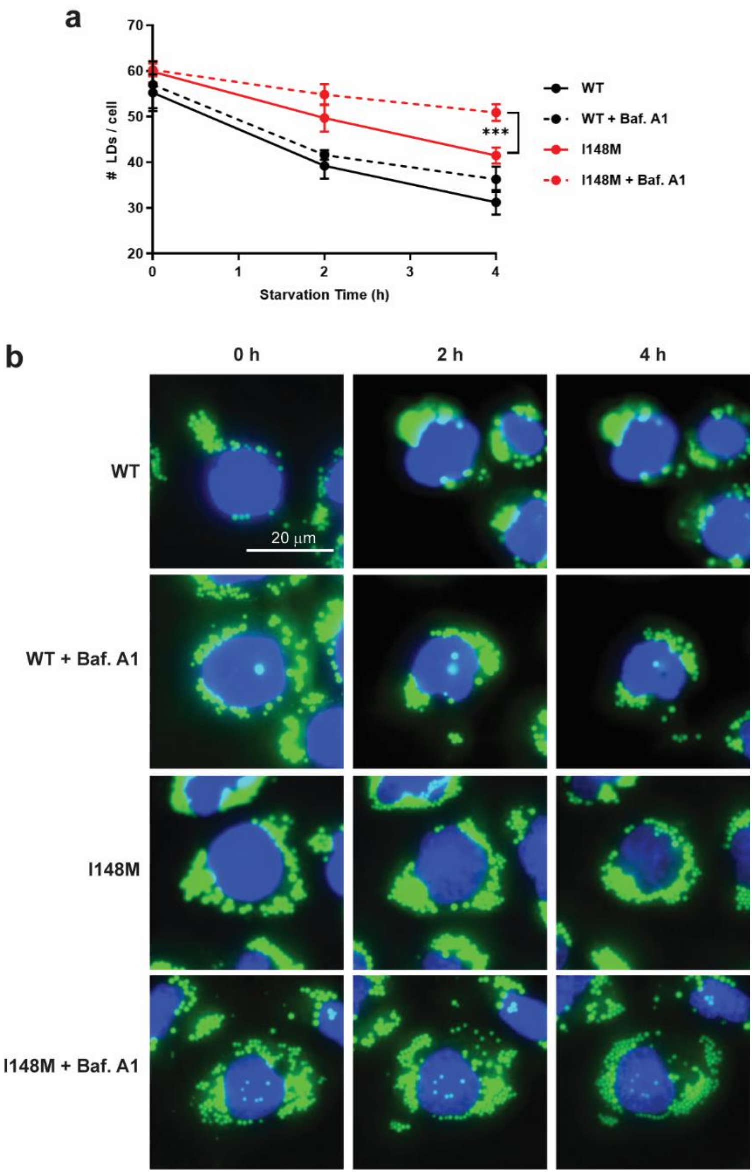


**Figure S6: I148M cells have a greater dependence on lipophagy than WT cells. a,** Hep3B cells were starved for four hours on a high-content imaging platform and lipid droplets were quantified at 0, 2, and 4 h. Some cells were treated with 100 nM Bafilomycin A1 to inhibit autophagy. LDs per cell were quantified. Analyzed by two-way Anova with Tukey's multiple comparisons test. 4 h: I148M-I148M + Baf. A1, adj. $p$ = 0.0002. **b,** Representative microscope images from the starvation time course. LDs and nuclei were stained with LipidTOX Green and Hoechst 33342 (blue), respectively. *** $p \leq 0.001$.

**Supplementary Table 1: Normalized *PNPLA3* I148I vs. I148M expression in primary human hepatocytes.**

| | | Relative Expression (dPCR values normalized to *TBP*) | | | |
|---|---|---|---|---|---|
| **Lot ID** | **Vendor** | **ANTZ9CG-I148M** | **ANTZ9CG-I148I** | **Total PNPLA3** | **Genotype** |
| **4056B** | Lonza | 0.000685408 | 0.475977972 | 0.48701422 | I148I |
| **4111C** | Lonza | 0 | 1.669507105 | 2.18944009 | I148I |
| **BVI** | BioIVT | 0 | 1.778876654 | 1.88584089 | I148I |
| **BXM** | BioIVT | 4.253581534 | 0 | 3.93356305 | I148M |
| **FZQ*** | BioIVT | 1 | 0 | 1 | I148M |
| **IXL** | BioIVT | 0.654735118 | 0.000485132 | 0.75487599 | I148M |
| **ONR** | BioIVT | 3.158128872 | 0 | 3.81124249 | I148M |

*All values relative to FZQ

### Footnote

† Created in BioRender. **Fig. S5a:** Sherman, D. (2024) BioRender.com/t23s872.
